# Single-Nucleus Transcriptomics Reveals Cell Type-Specific Remodeling and Epilepsy-Associated Microglia

**DOI:** 10.64898/2026.03.04.709339

**Authors:** Victoria Ho, Ruth Tjondropurnomo, Jennifer Nguyen, Eszter Balkó, Samantha Depew, Xing Chen, Radhika Singh, J Edward Van Veen, Bence Rácz, Peyman Golshani

## Abstract

Mesial temporal lobe epilepsy (TLE) is the most common form of acquired epilepsy involving the hippocampus and is a frequent sequelae of head trauma. TLE is associated with refractory seizures and significant cognitive deficits. Yet, the gene expression patterns and cell types driving epileptogenesis and the associated cognitive deficits are poorly understood. To address this, we performed single nucleus RNA sequencing on hippocampal tissue from mice at 3 and 6 weeks following pilocarpine-induced status epilepticus, a robust model of TLE. At these early timepoints, epilepsy samples showed reductions in specific Cck and Lamp5-Lhx6 interneuron subclusters, alongside increases in Cajal-Retzius cells, dentate granule (DG) cell precursors, and a mature DG cell subcluster. Among glia, an astrocyte subcluster and a markedly expanded microglia sublcuster were increased. We term this microglia population epilepsy-associated microglia (EAM). The transcriptomic profile of EAM partially overlaps with microglia described in models of Alzheimer’s disease and traumatic brain injury, with enrichment of genes including *Myo1e* and *Igf1*. EAM display amoeboid morphology, can be found in dense clumps around pyramidal and granule cell body layers, and exhibit enlarged vesicles and mitochondria on electron microscopy. Cell-cell interaction analysis predict that DG cells are the main interaction partners of EAM. This dataset recapitulates known cellular alterations in TLE while defining their underlying transcriptomic programs, enabling mechanistic dissection of the key processes driving epileptogenesis.

## Introduction

Epileptogenesis is the process by which brain tissue becomes capable of generating spontaneous seizures^1^. In acquired epilepsy, an acute insult precipitates epileptogenesis, initiating molecular and cellular events that lead to a dysfunctional network that is chronically prone to seizures. The latency between the insult and the clinical onset of epilepsy can be months to years in humans, presenting a window of opportunity to intervene on the epileptogenic processes^2^. However, despite intensive efforts, no disease-modifying therapies are currently available. A major obstacle to the development of anti-epileptogenic therapies is an incomplete understanding of the key processes driving epileptogenesis.

Mesial temporal lobe epilepsy (TLE) is the most common form of acquired epilepsy in adults and is frequently refractory to anti-seizure medications^3^. Epileptogenesis in TLE is characterized by pathological remodeling of hippocampal circuits. In both patients and experimental models, the remodeling includes selective neuron loss, aberrant neurogenesis leading to ectopic granule cells, gliosis, and axonal sprouting, which result in long-lasting alterations in hippocampal circuitry^4^. In addition to generating spontaneous, recurrent seizures, epilepsy-associated changes in hippocampal circuits result in less precise and stable hippocampal activity patterns^5,6^, resulting in domain-specific cognitive deficits. In TLE, the characteristic deficit is episodic memory impairment^7^. Prior transcriptomic studies have identified numerous gene expression changes occurring at various time points during epileptogenesis. Acute periods following injury are associated with the largest changes in gene expression, whereas later periods show changes in genes with a greater diversity of cellular function^8^. However, the significance of gene expression changes are difficult to interpret without knowing the cell type in which they are occurring, and a clear conceptual framework for key processes has been out of reach.

The hippocampus comprises many neuronal and glial cell types with incompletely understood roles in generating the epileptic network. Here, we take an unbiased approach using single nucleus RNA sequencing (snRNAseq) to dissect this cellular heterogeneity and define cell type-specific transcriptional programs engaged in hippocampal epileptogenesis. We use a well-established model of TLE in which pilocarpine is used to induce an episode of status epilepticus (SE), which acts as a precipitating injury to trigger epileptogenesis. We performed tissue collection at 3 and 6 weeks post-SE based on our previous work which showed that place cell activity is normal at 3 weeks but becomes less precise and less stable at 6 weeks, demonstrating active, functionally consequential network remodeling during this time window^9^. We found dramatic changes in gene expression in a number of cell types in epilepsy mice. Notable changes were increases in abundance of neuroblasts and immature dentate granule cells in the TLE model, reflecting a pathological form of adult neurogenesis. We also found subclusters within interneuron subtypes that were decreased in the TLE model, demonstrating selective vulnerability within specific cell types that can now be transcriptionally characterized. Lastly, we identified an epilepsy-associated subtype of microglia (EAM) that are abundant in epilepsy and nearly completely absent from controls. These results provide a high-resolution atlas of hippocampal remodeling in TLE and establish a framework for understanding how cell type-specific molecular changes contribute to the development of chronic epilepsy and the associated cognitive dysfunction.

## Results

### A single-nucleus transcriptomic atlas of the hippocampus during epileptogenesis

To model TLE, wild-type mice were given intraperitoneal injections of pilocarpine to induce SE, thereby initiating epileptogenesis. To minimize the confounding transcriptomic effects of acute seizures, all mice were monitored prior to tissue collection with electroencephalography (EEG) to ensure a 12-hour period of seizure freedom. Bilateral dorsal hippocampi were collected from each mouse at 3 and 6 weeks post-SE for single nucleus RNA sequencing (Figure 1A). A total of 16 epileptic and 15 control animals were sequenced, which yielded 151,034 nuclei after applying quality control thresholds. Reads were processed using Seurat, clustered, and annotated into different cell types (Figure 1B-E). Cell-type annotation was based on canonical marker gene expression as well as label transfer using reference datasets^10,11^. All major cell types were represented in our dataset. Additionally, the large number of nuclei sequenced enabled identification of less abundant populations including mossy cells (*Rgs12*^+^/ *Necab1*^+^/ *Drd2*^+^) and Cajal-Retzius neurons (CR; *Reln*^+^/*Trp73*^+^/*Lhx1*^+^/*Slc17a7*-*Slc17a6*^mixed^), as well as cell types in the neurogenesis lineage: radial glia-like (RGL) cells (*Gfap*^+^/*Sox9*^+^/*Notch1*^+^/*Hopx*^+^/*S100b*^-^), neuroblasts (*Eomes*^high^/*Sox11*^+^/*Dcx*^+^), and immature dentate granule cells (*Eomes*^low^/*Sox11*^+^/*Dcx*^+^) (Figures 1B and C, S1, S2, S3A and B).

**Figure 1.**
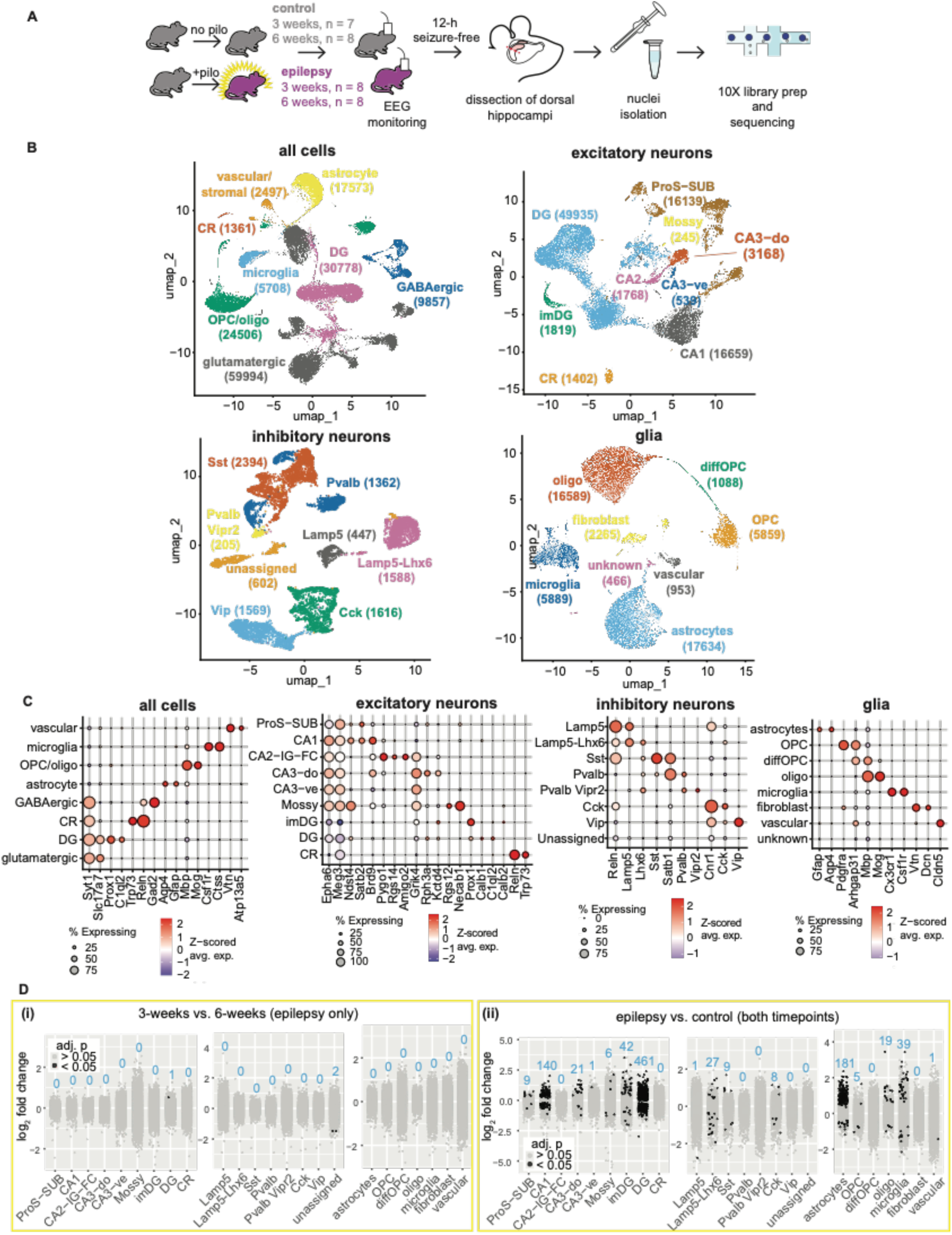
Single-nucleus RNA sequencing reveals the cellular landscape of the hippocampus during epileptogenesis. (A) Experimental overview. (B) UMAP visualization of 151,034 single nucleus RNA profiles colored by cluster. The number of nuclei in each cluster is shown in brackets. (C) Dot plot showing expression of select marker genes. Dot size represents the percentage of cells in the cluster expressing the gene. Dot color represents the scaled average expression (z-scored across clusters). (D) Pseudobulk differential gene expression comparing (i) epilepsy samples at 3 versus 6 weeks post-SE and (ii) epilepsy versus control samples with both timepoints combined. Black and gray dots represent genes with adjusted p <0.05 and >0.05, respectively. Blue numbers indicate the number of genes with adjusted p-value <0.05. OPC, oligodendrocyte progenitor cell; diffOPC, differentiating oligodendrocyte progenitor cell; DG, dentate granule cells; imDG, immature dentate granule cells; CR, Cajal-Retzius cells; ProS, pro-subiculum; SUB, subiculum. [High-resolution image available as separate file.]

To identify differentially expressed genes (DEGs), single nuclei counts were aggregated for pseudobulk analysis. We first compared gene expression in epilepsy samples between the 3- and 6-week post-SE timepoints, given the deterioration in place cell function seen during this period in our prior study^9^. However, there were very few genes that were differentially expressed between the two timepoints (Figure 1D(i); Extended Data Table 1). Interaction testing also identified essentially no genes with significant time-dependent epilepsy effects across all cell types, indicating that transcriptional changes in epilepsy were largely stable between 3 and 6 weeks (Extended Data Table 1). On the other hand, there were numerous significant DEGs when comparing epilepsy to control samples with both timepoints combined (Figure 1D(ii); Extended Data Table 1). These findings suggest that gene expression is altered across many cell types in epilepsy but remain relatively stable between the 3- and 6-week timepoints, indicating that the functional changes observed in place cells may be driven by post-transcriptional mechanisms or by changes to hippocampal inputs^12^.

### Cell type composition is altered in epilepsy

To evaluate changes in the abundance of different cell types, we calculated their proportions within each sample as a fraction of the total nuclei sequenced from that sample (Extended Data Table 2). We then examined which factors influenced the proportions. Multivariate analysis revealed a robust effect of pilocarpine treatment on overall cellular composition, while neither timepoint (3 vs. 6 weeks post-SE) nor its interaction with treatment significantly influenced cell type proportions (Extended Data Table 3). These findings are consistent with the abundant number of DEGs we found between epilepsy and control, and the paucity of DEGs when comparing 3- and 6-week epilepsy samples.

We compared cell type proportions between control and epilepsy to identify changes associated with epileptogenesis. The proportions of several cell types capable of local proliferation within the hippocampus were increased, including oligodendrocytes (fold change, FC 1.86, false discovery rate, FDR 0.017), the immature dentate granule cell (imDG) cluster (FC 2.83, FDR 0.017), and microglia (FC 3.70; FDR 0.038) (Figure 2A,B; Table 4). We also found an increase in Cajal-Retzius (CR) cells (FC 1.51, FDR 0.039), corroborating findings in human TLE where increased levels are presumably due to enhanced persistence of CR neurons into adulthood^13^. While astrocyte proliferation has been found to occur following induced SE and provoked seizures^14,15^, we did not see an increase in our samples, perhaps due to the exclusion of acute seizures prior to tissue collection.

**Figure 2.**
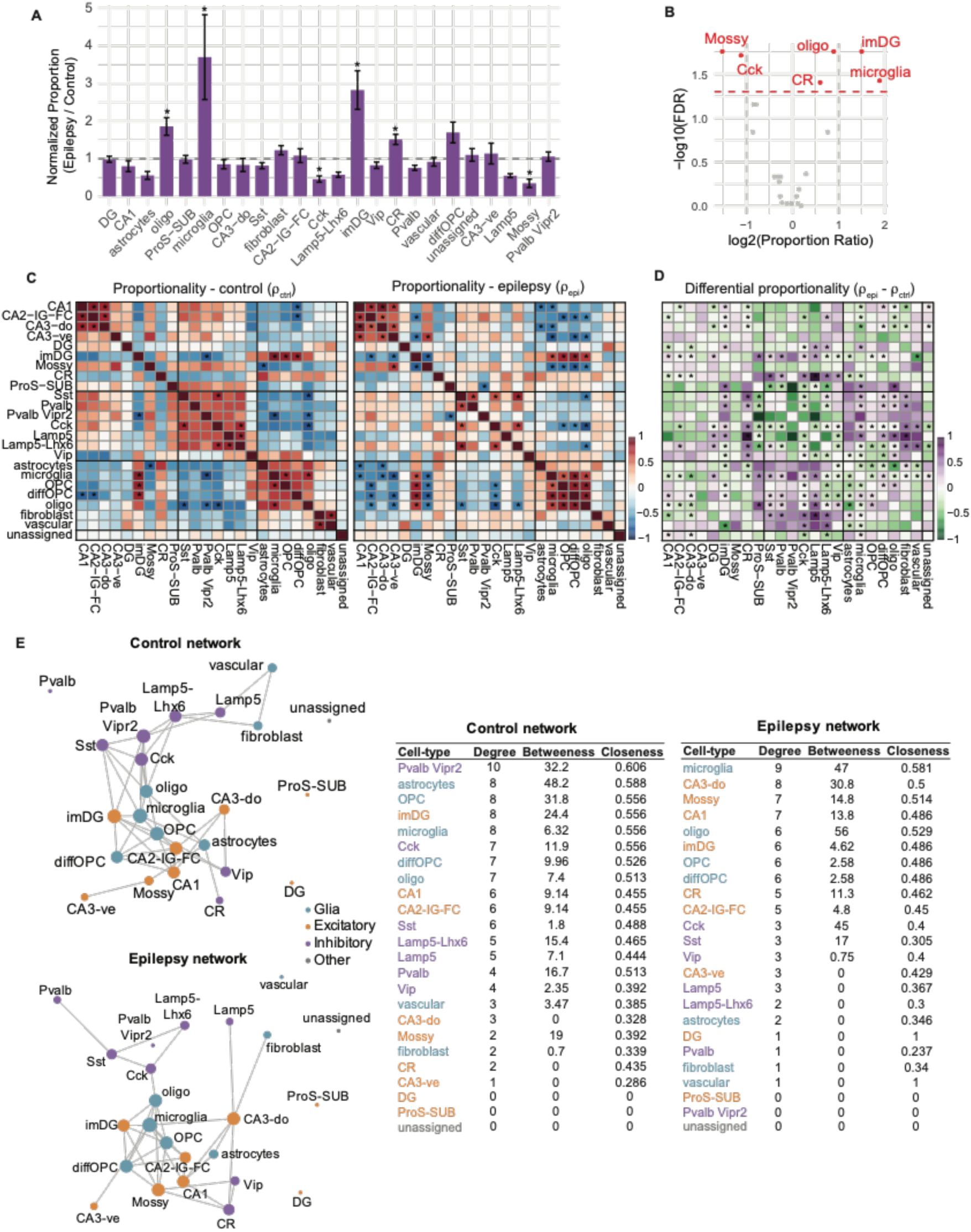
Changes in cellular composition in epilepsy. The abundance of different cell types was quantified as proportion of total cells sequenced within a sample. (A) Bar graphs show relative cell type proportions in epilepsy normalized to control (epilepsy/control fold change). Error bars show standard error of the mean. * p <0.05. (B) Volcano plot showing magnitude of proportional change (log_#_ fold change) versus statistical significance for each cell type. (C) Heatmaps showing how cell types vary together across samples within each condition (control vs. epilepsy), i.e. pairwise proportionality (𝜌). Positive values indicate that two cell types tend to increase or decrease together across animals, while negative values indicate that when one increases, the other decreases. At FDR ≤ 0.05, significant pairs were those with |𝜌| ≥ 0.71 (control) and |𝜌| ≥ 0.60 (epilepsy). (D) Heatmap showing how proportionality changes in epilepsy, i.e. differential proportionality (𝛥𝜌 = 𝜌*_epilepsy_*− 𝜌*_control_*). Positive values indicate stronger coordination between two cell types in epilepsy compared to control; negative values indicate weaker coordination. Asterisks mark pairs with FDR ≤ 0.05. (E) Network diagrams summarizing coordinated changes in cell type abundance, inferred using CLR-transformed and 𝑧-scored cell type proportions. Nodes represent cell-types and edges represent partial correlations. [High-resolution image available as separate file.]

Cholecystokinin neurons (Cck; FC 0.46, FDR 0.019) and mossy cells (FC 0.35, FDR 0.017) were significantly decreased (Figure 2A,B; Table. 4). Overall, these findings recapitulate prior findings in TLE of interneuron and mossy cell loss, dentate granule cell proliferation, and gliosis^4^.

We next examined whether changes in epilepsy extend beyond individual cell types to coordinated shifts in cellular composition. We estimated pairwise proportionality (𝜌) for all cell types within each sample and found co-variation among several pairs (Figure 2C).

Differential proportionality analysis (Δ𝜌 = 𝜌*_epi_* – 𝜌*_ctrl_*) revealed increased coupling between glial and interneuron populations in epilepsy (Figure 2D). Specifically, glia-interneuron pairs showed the strongest and most pervasive increase in proportional co-variation with 20 significant positive pairs and the largest overall increase in coordinated variation (∑Δ𝜌 = 5.83) (Extended Data Table 5). In contrast, glia-excitatory neuron pairs showed an opposite trend with net negative shift (∑Δ𝜌 = −1.31). Interneuron-interneuron pairs also tended to decrease in proportional co-variation, with 2 positive and 7 negative significant pairs and a net negative shift (∑Δ𝜌 = −1.20) (Extended Data Table 5). We next applied network inference analysis to identify cell types that may serve as central coordinators (hubs) of changes in cell abundance. This analysis identified microglia as a central hub cell type, with the largest number of edges (which connect cell types with correlated proportions) and highest closeness (i.e. inverse of the average distance to all other nodes). This network position suggests that microglia expansion may contribute to or reflect broader shifts in cellular composition. Notably, many edges connecting interneuron populations that were present in controls were absent in epilepsy, consistent with decoupled alterations in interneuron abundance (Figure 2E).

### Predicted cell–cell signaling is altered in epilepsy

To further investigate whether specific cell types drive changes during epileptogenesis, we used CellChat to predict incoming and outgoing interactions for each cell type. Interaction strength (weight) was first determined with replicates pooled for each condition. This approach suggested that microglia have the largest changes in both incoming and outgoing interactions (Figure 3A).

**Figure 3.**
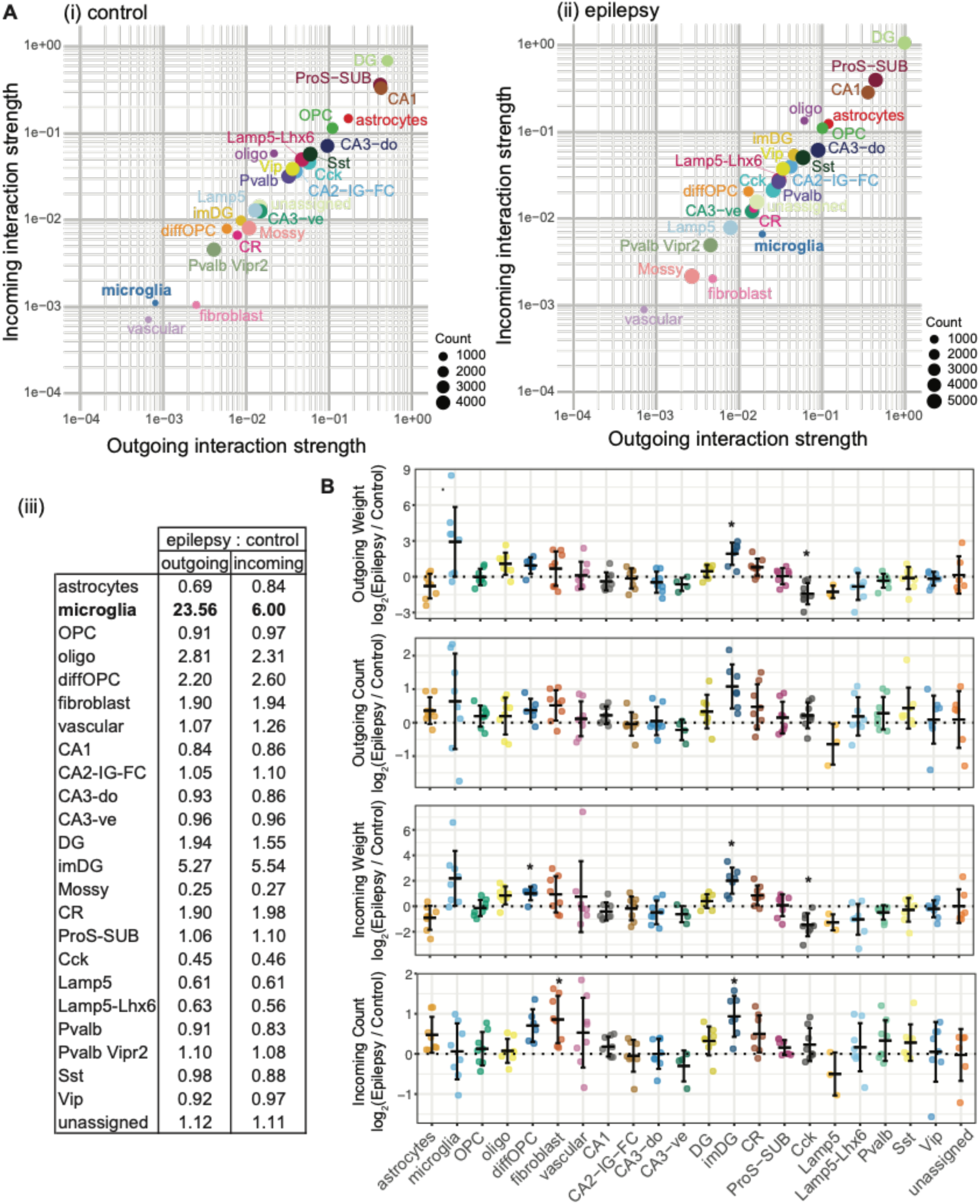
Predicted cell–cell communication is altered in epilepsy. (A) Cell-cell interactions predicted with samples pooled for each state. Incoming vs. outgoing interaction strengths plotted on log-scale axes for (i) control and (ii) epilepsy. (iii) Table shows interactions strengths in epilepsy normalized to control. (B) Cell-cell interactions predicted for each sample. The weight (strength) and count (number) of incoming and outgoing interactions were determined separately for each sample. Within each batch, log_#_ epilepsy/control values were computed and tested against zero using a Student’s t-test. Bars represent mean ± s.d. Significance markers for BH adjusted p-values: * p <0.05. See Extended Data Table 7. [High-resolution image available as separate file.]

Because pooling replicates may result in spurious predictions (e.g. ligands and receptors not present in the same samples) or over-weighted predictions (e.g. ligands and receptors represented at inflated levels), we repeated the predictions on a per-sample basis. At the per-sample level, neither the strength nor the number of microglial interactions differed significantly from zero when expressed as log_#_(epilepsy/control) ratios (Figure 3B; Extended Data Table 7). However, examination of individual samples revealed substantial variability in microglial interactions, with several showing marked increases, indicating heterogeneous microglial engagement across animals.

In contrast to microglia, log_#_(epilepsy/control) ratios for imDG were significantly altered across several interaction metrics, suggesting more consistent and robust changes in cell–cell interactions for this population.

### Cell-type subpopulations are differentially altered in epilepsy

To better resolve cellular heterogeneity and explore state-specific transcriptional profiles, each cell type was subsetted, re-integrated and re-clustered into subclusters. Subcluster proportions were calculated as fractions within each parent cluster to minimize biases in nuclei extraction that may differentially affect one cell type over another. This led to the identification of several notable changes in subcluster composition (Figure 4, Extended Data Table 6).

**Figure 4.**
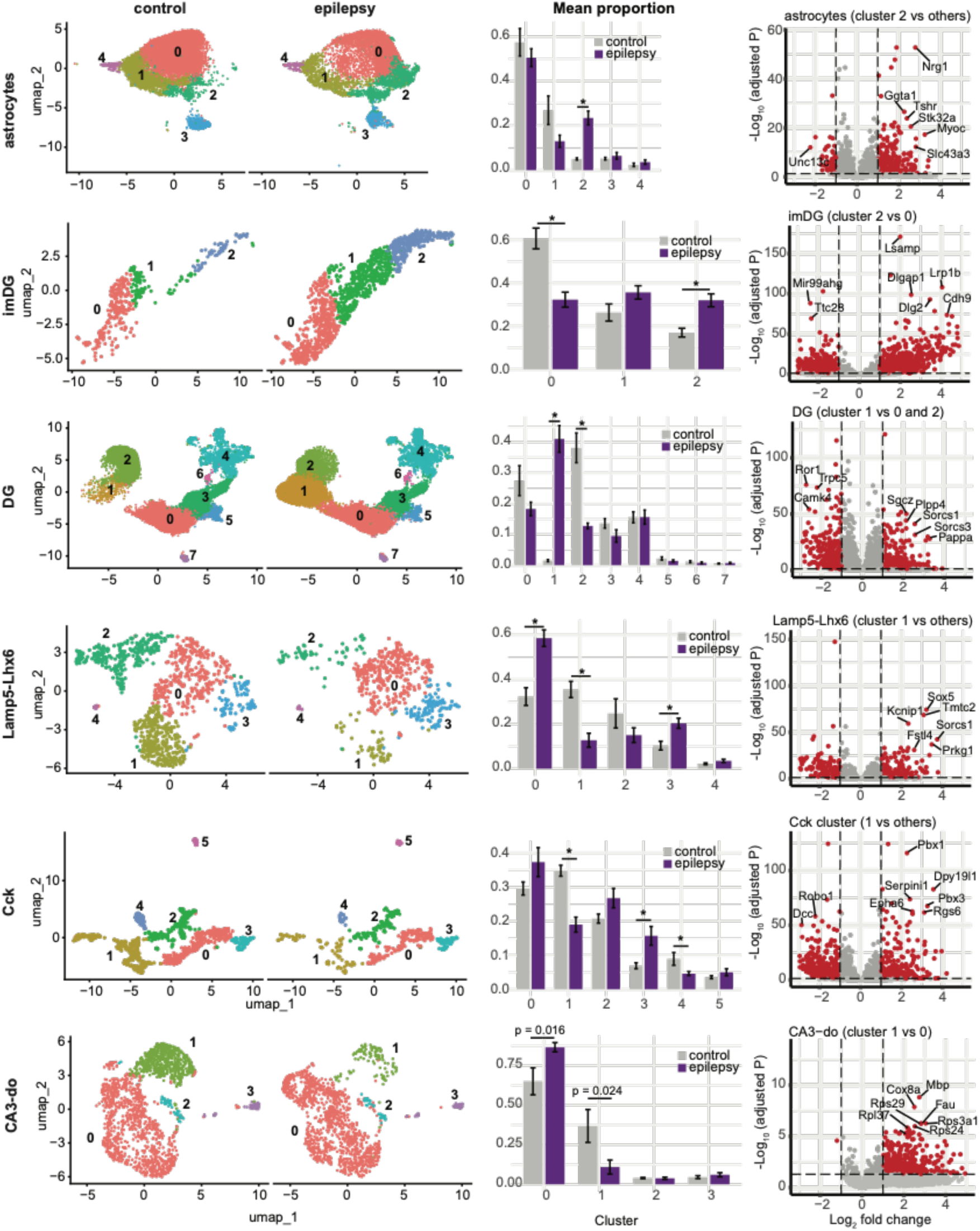
Epileptogenesis alters transcriptional subpopulations within major hippocampal cell types. Left: UMAPs of re-clustered cell types with significant changes in subcluster proportions. The plots are split by state to show control and epilepsy separately. Middle: Changes in subcluster proportions. Asterisks denote subclusters with global FDR <0.05. Right: Volcano plots of DEGs (also see Extended Data Table 6). See Figure S4 for UMAPs of cell types with non-significant changes in subcluster proportions. [High-resolution image available as separate file.]

Among astrocytes, the astrocytes_2 subcluster was significantly expanded in epilepsy (Figure 4). The imDG cluster increased across all subclusters (Figure 4), consistent with its overall expansion (Figure 2A,B). Although imDG_0 comprised a smaller fraction of the imDG cluster, its absolute abundance was increased in epilepsy (304 nuclei vs 150 nuclei). Examination of canonical cell type markers and reference-based annotation indicates that the imDG cluster comprises neuroblasts in addition to immature dentate granule cells (Figure S2)^11^.

Within the dentate granule (DG) cell cluster, there was a large increase in the proportion of the DG_1 subcluster in epilepsy and a corresponding decrease in the DG_2 subcluster.

Although we detected increased neurogenesis, DG_1 makes up about 40% of the parent DG cluster and is hence too abundant to be solely accounted for by adult neurogenesis. Instead, trajectory inference suggests that DG_1 represents a terminal state derived from DG_2 (Figure S3), consistent with a shift in DG cell state.

Among inhibitory neurons, specific Lamp5-Lhx6 and Cck subclusters were reduced in epilepsy, indicating selective vulnerability (Figure 4; Extended Data Table 6). A CA3-do subcluster also declined, but the difference was not significant after multiple testing correction. While neuronal loss is a hallmark of TLE, these findings suggest that vulnerability is not uniform but instead is concentrated in discrete transcriptional subpopulations. The resulting epileptic networks reflect both loss of vulnerable subpopulations and preservation of others.

Most strikingly, a distinct microglial subcluster was highly enriched in epilepsy but nearly completely absent from controls (Figure 5A,B), highlighting a robust and disease-associated glial response.

**Figure 5.**
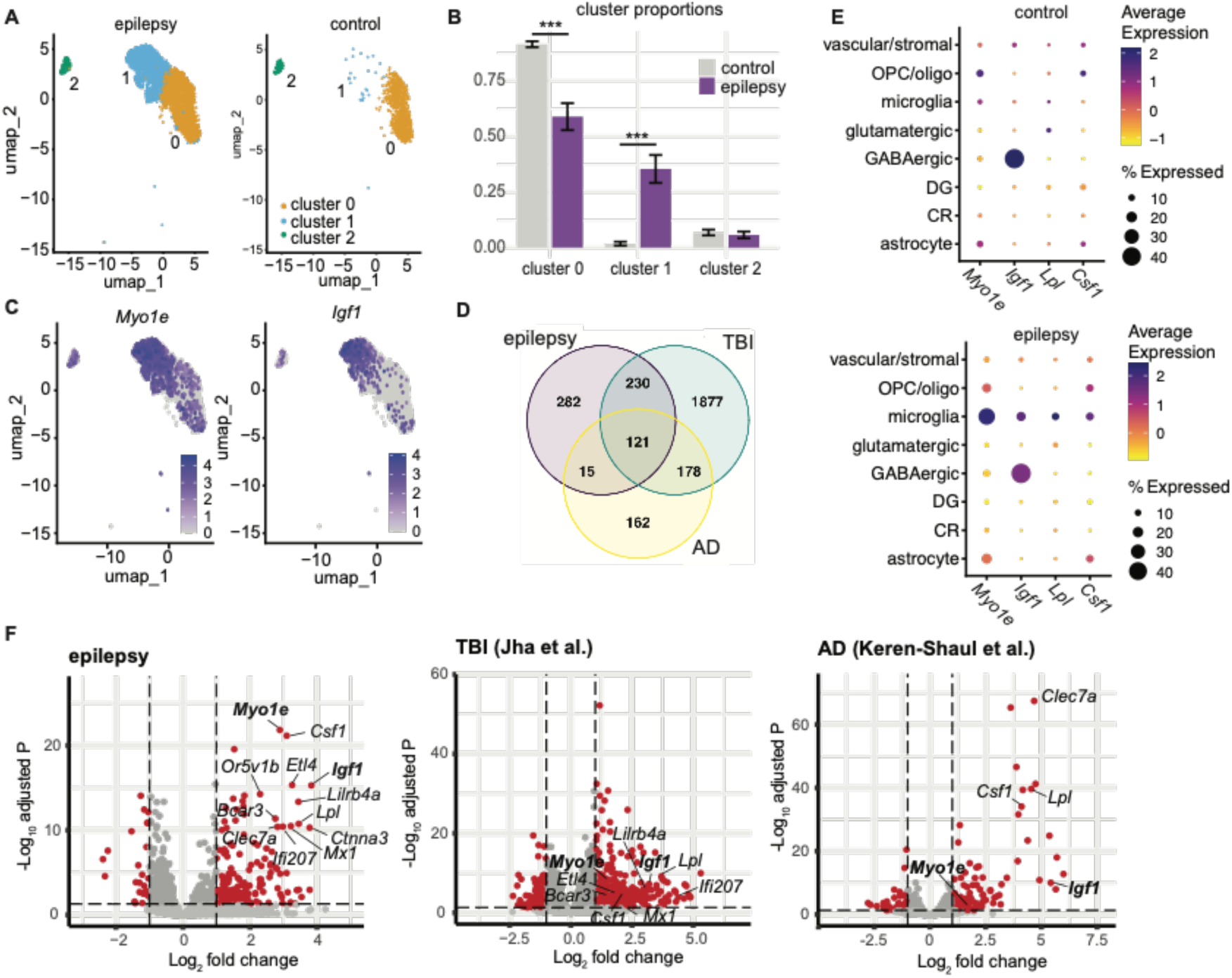
Identification of an epilepsy-associated microglia subcluster. (A) UMAP of re-clustered microglia split by state to show control and epilepsy separately. (B) Changes in subcluster proportions. Error bars show SEM. Asterisks denote FDR <0.001. (C) FeaturePlot showing expression of two EAM-enriched genes, Myo1e and Igf1, in microglia subclusters. (D) Venn diagram showing overlap of differentially-expressed genes (adjusted P <0.05) in disease-associated microglia. There were 121 genes shared across all three conditions, representing a significant overlap (fold enrichment = 1.34, 𝑃 = 1.14166 × 10^-4^, SuperExactTest, background n = 2,865). (E) Dot plot showing average expression of EAM-enriched genes in different cell types in control (top) and epilepsy (bottom). (F) Volcano plot of genes that are differentially expressed in disease-associated subclusters of microglia in epilepsy (left), TBI (middle)^17^, and AD (right)^18^. Labels show genes enriched in EAM with 𝐿𝑜𝑔_#_fold change >2 and −𝐿𝑜𝑔_01_adjusted P >10. [High-resolution image available as separate file.]

### A distinct microglial population emerges in epilepsy

Subsetting and re-clustering of microglia resulted in two main clusters – a cluster present in both epilepsy and control (microglia_0) and a cluster abundant in the epilepsy samples but nearly completely absent from controls (microglia_1) (Figure 5). We refer to the latter subcluster as epilepsy-associated microglia (EAM). EAM express many canonical microglia markers, including *Cx3cr1*, *Aif1*, and *P2ry12* (Figure S5). In addition to microglia_0 and EAM, there was a small subcluster that expressed markers for border-associated macrophages (Figure S6B). To evaluate the robustness of EAM, we repeated integration and clustering, leaving out one batch of samples at a time, which consistently identified an epilepsy-enriched subcluster in each iteration (Figure S7). We also confirmed that dissociation-related artifacts were minimal in our microglia clusters by checking immediate early gene expression levels (Figure S8)^16^.

Disease-associated microglia have been identified in models of other neurological conditions, including two that are risk factors for developing TLE: Alzheimer’s disease (AD) and traumatic brain injury (TBI). Comparison of DEGs in the disease-associated microglia of all three models identified 121 genes that were present in all three conditions (EAM, TBI, and AD) (Extended Data Table 8). The triple overlap is significant (𝑃 = 1.14166 × 10^-4^) (Figure 5D) ^17,18^.

To select suitable markers for further characterization of EAM, we examined the expression patterns of the genes that were most enriched and significant in EAM (Figure 5F). Among these, we focused on genes that were abundant in EAM and relatively absent from microglia_0 (Figure 5C), and also had low expression in other cell types (Figure 5E). Our last criteria was that the genes should be enriched in disease-associated microglia from AD and TBI models as well (Figure 5F)^17,18^. This led to the selection of *Myo1e* and *Igf1*.

Microglia-derived *Igf1* has been reported to act as a trophic factor in maintaining neuronal survival during postnatal development of layer V cortical neurons^19^. *Myo1e* has been studied in peripheral immune cells and found to be involved in phagocytosis^20^.

### Histological validation of EAM expansion in hippocampal tissue

Immunohistochemistry (IHC) combined with fluorescence in situ hybridization (FISH) confirmed the presence of *Igf1-/Myo1e-* and *Igf1+/Myo1e+* populations of microglia, the latter corresponding to EAM (Figure 6A). EAM comprised up to a third of the microglia population (Figure 6B), similar to their proportion in the snRNAseq data. We also detected elevated levels of MYO1E protein in epilepsy samples (Figure 6G,H). We were unable to find an IGF1 antibody that worked well for immunostaining.

**Figure 6.**
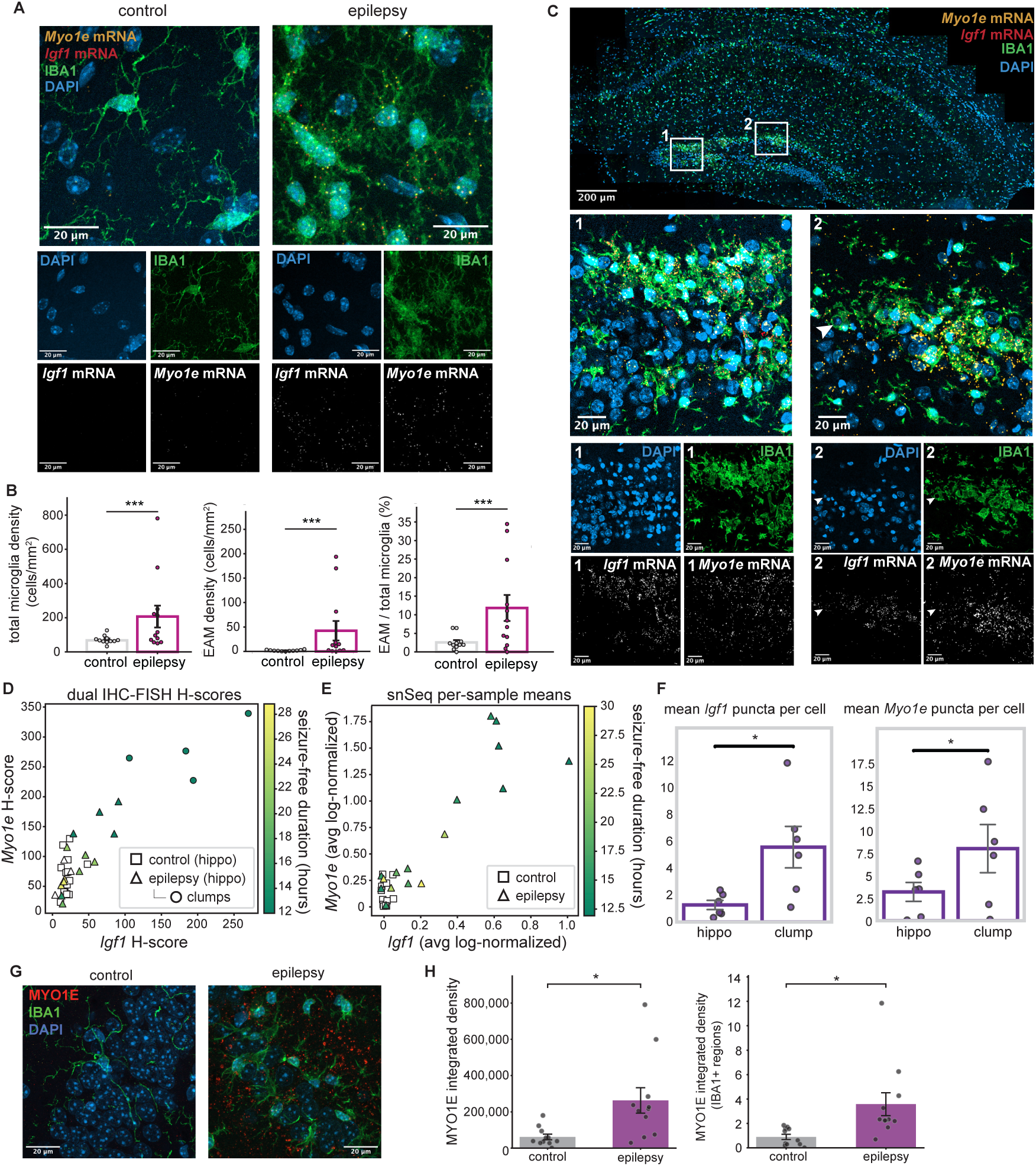
Histological validation of EAM expansion and marker expression. (A) Representative IHC-FISH images of hippocampal tissue from control (left panel) and epilepsy (right panel). Large square shows merged image. Yellow, Myo1e mRNA; red, Igf1 mRNA; green, IBA1 protein; blue, DAPI nuclear stain. (B) Quantification of total microglia and EAM density from IHC-FISH images. Each point represents a biological replicate. (C) IHC-FISH images of hippocampal tissue from epilepsy. Large rectangle shows a tiled image with enlarged insets shown below. Yellow, Myo1e mRNA; red, Igf1 mRNA; green, IBA1 protein; blue, DAPI nuclear stain. (D) Plot of Myo1e vs. Igf1 H-scores quantified from hippocampal sections from IHC-FISH images. Markers are colored by seizure-free duration. Each square and triangle represents a biological replicate from control and epilepsy, respectively. Each circle represents a clump of microglia from an epilepsy sample. (E) Plot of average log-normalized Myo1e vs. Igf1 levels in the microglia cluster from snRNAseq data. Markers are colored by seizure-free duration. Each square and triangle represents a biological replicate from control and epilepsy, respectively. (F) Quantification of Myo1e and Igf1 mRNA puncta per IBA+ cell in entire hippocampi and in microglia clumps. Each point represents a biological replicate. (G) Representative IHC images of hippocampal tissue from control (left panel) and epilepsy (right panel). Red, MYO1E protein; green IBA1 protein; blue, DAPI. (H) Quantification of (G). Left, integrated density of MYO1E signal in entire hippocampal regions. Right, integrated density of MYO1E signal that overlaps with IBA1 signal. Each point represents a biological replicate. In all panels, error bars indicate SEM. Significance markers: * p <0.05, ** p <0.01, *** p <0.001. [High-resolution image available as separate file.]

Microgliosis and EAM were present throughout hippocampal CA1-3 and DG subregions of epilepsy samples (Figure S9A, Extended Data Table 9). In both total microglia and EAM density measurements, epilepsy samples exhibited right-skewed distributions driven by a subset of animals with very high densities (Figure 6B). Consistent with this, variability was markedly greater in epilepsy than in controls for total microglia density (SD*_epilepsy_* = 221.46, SD*_control_* = 23.99, variance ratio = 85) as well as for EAM density (SD*_epilepsy_* = 69.06, SD*_control_* = 1.38, variance ratio = 2519). Variance was compared using the Fligner-Killeen test and revealed significantly greater variability in epilepsy for both total microglia density and EAM density (total microglia: p = 0.0037; EAM: p = 6.6 x 10^-4^).

To assess the effect of disease state and account for potential confounders, we performed an ANCOVA on total microglia density with state (epilepsy vs. control) as the primary factor, and sex, weeks post-SE, and seizure-free duration as covariates. Epilepsy was significantly associated with increased total microglial density (𝐹(1,15) = 21.59, 𝑝 = 3.16 × 10^-4^) (Figure 6B). Notably, longer seizure-free duration independently predicted lower microglial density (𝐹(1,15) = 14.21, 𝑝 = 0.00186), indicating that more recent seizures were associated with greater microgliosis. Sex (p=0.517) and weeks post-SE (p=0.148) were not significant predictors (Extended Data Table 10; Figure S9B, C). A similar ANCOVA for EAM density revealed that epilepsy strongly predicted increased EAM density (𝐹(1,15) = 26.25, 𝑝 = 1.25 × 10^-4^). Seizure-free duration again emerged as an independent negative predictor (𝐹(1,15) = 18.73, 𝑝 = 5.98 × 10^-4^), while sex (p=0.577) and weeks post-SE (p=0.076) were not significantly correlated (Extended Data Table 11; Figure S9B, C).

The relationship between seizure-free duration and microglia abundance was further assessed with Spearman’s rank correlation, which identified an inverse correlation between seizure-free duration and both total microglia density (𝜌 = −0.83, p = 0.005) and EAM density (𝜌 = −0.917, p = 0.000507) (Figure S9D). Longer seizure-free intervals were also associated with lower *Igf1* (Spearman rho =-0.683, p = 0.0424) and *Myo1e* (Spearman 𝜌 = −0.700, p = 0.0358) expression levels, as quantified with H-scores (Figure 6D, S9E). The snRNAseq data showed similar negative trends with seizure-free duration, although these relationships did not reach statistical significance (Figure 6E, S10), perhaps due to the compositional nature of snRNAseq data and technical limitations such as transcript dropout.

EAM were occasionally seen in clumps where they had indistinguishable cell boundaries and appeared to infiltrate pyramidal and granule cell body layers (Figure 6B, S11). EAM within clumps tended to have higher levels of *Igf1* and *Myo1e* puncta (Figure 6E, F).

### EAM represent an activated microglial state

To characterize morphological differences in microglia, we quantified their ramification index (RI), which is calculated as the territorial volume of a microglial cell divided by its cellular volume. Higher ramification indices indicate a more branched morphology while lower indices indicate a more compact morphology (Figure 7A). EAM (*Igf1+; Myo1e+*; RI = 5.518 ± 0.398) had significantly lower RIs than *Igf1-; Myo1e-* microglia in control (15.88 ± 0.961), but not *Igf1-; Myo1e-* microglia in epilepsy (8.31 ± 1.279) (Figure 7B). Hence, in epilepsy mice, the *Igf1-; Myo1e-* microglia (corresponding to microglia_0) were morphologically more similar to the *Igf1+; Myo1e+* population (microglia_1 / EAM) within epilepsy samples than the *Igf1-; Myo1e-* microglia in control animals. This indicates that, despite occupying a similar transcriptomic space to control microglia_0 (low *Igf1; Myo1e* expression), microglia_0 in epilepsy adopt a morphological phenotype more similar to EAM.

**Figure 7.**
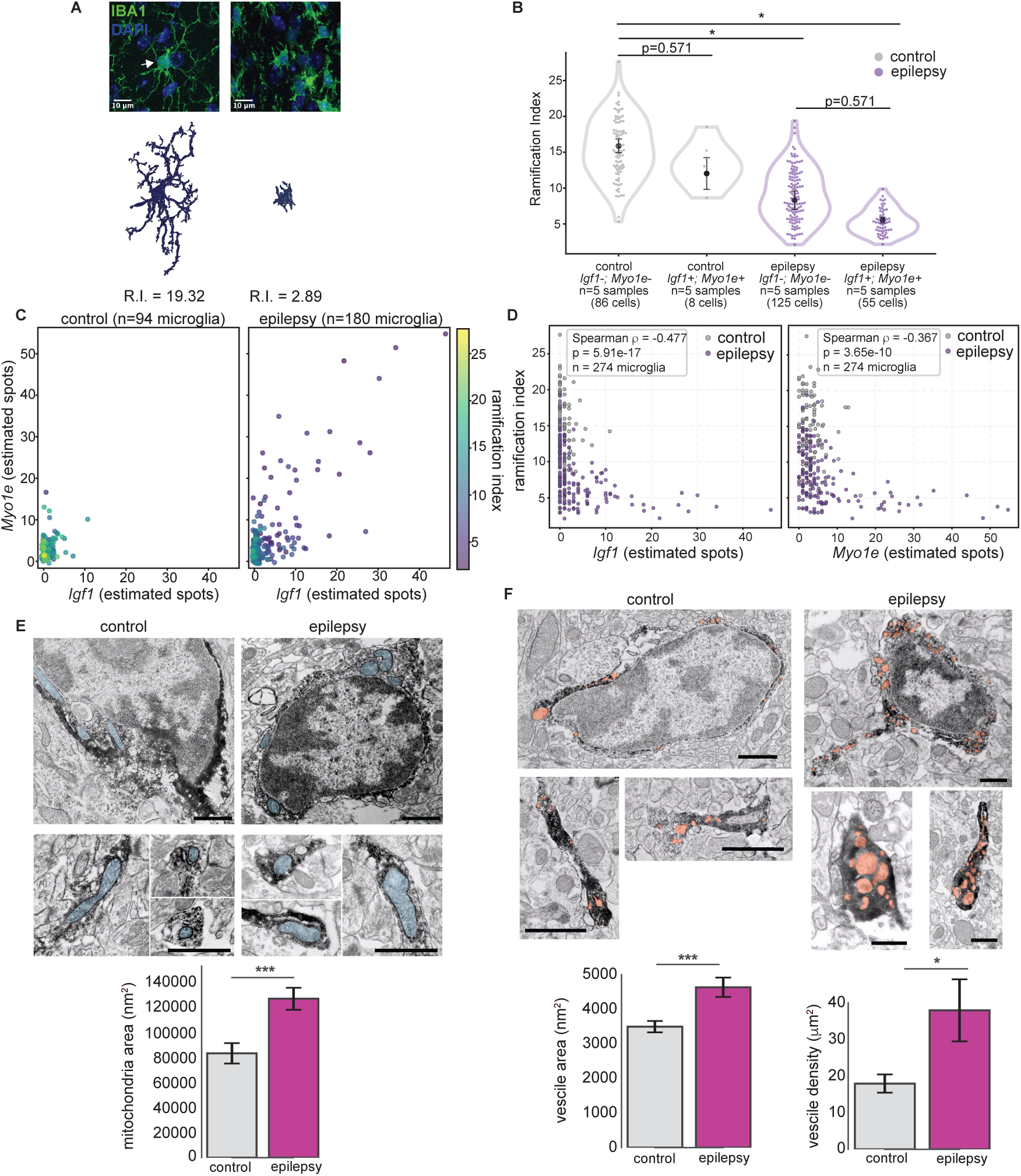
EAM exhibit features of activated microglia. (A) Examples of high and low RI. Green, IBA1 protein; blue, DAPI nuclear stain. (B) Violin and swarm plot of per-cell RI. Error bars show mean ± SEM of per-sample RI. P-values are from two-sided Mann-Whitney tests on per-sample means with Holm adjustment. Number of biological replicates shown on the x-axis. (C) Quantification of Igf1 vs. Myo1e expression from IHC-FISH images. Each marker represents a cell and is colored by RI. (D) Correlation between RI and Igf1 (left) or Myo1e (right) expression levels. Each point represents a microglial cell, colored by disease state. Pooled Spearman correlation coefficient (𝜌), p-value, and total number of cells are indicated. (E) Representative electron microscopy images from Iba1 immunolabeled tissue with vesicles pseudo-colored in orange. Scale bar = 1000 nm. Bar graphs show quantification performed on 10-20 sections per animal (n = 3 animals per condition). (F) Representative electron microscopy images from Iba1 immunolabeled tissue with mitochondria pseudo-colored in blue. Scale bar = 1000 nm. Bar graphs show quantification performed on 10-20 sections per animal (n = 3 animals per condition). Significance markers: * p <0.05, ** p <0.01, *** p <0.001. [High-resolution image available as separate file.]

Subcellular organelle function within microglia is altered in pathological states and are closely linked to microglial function^21^. To investigate how mitochondria and lysosomes are altered in epilepsy, we examined hippocampal tissue from epilepsy and control animals at 6 weeks post-treatment using electron microscopy. In the epilepsy samples, membranous vesicle-like structures – likely representing lysosomes and endosomes – were larger and approximately 2-fold more numerous than in controls (Figure 7E), indicating heightened engagement of the endo-lysosomal system. Mitochondria also appeared enlarged in epilepsy samples (Figure 7F), a phenotype that has similarly been observed following Lipopolysaccharide challenge^22^.

### EAM are predicted to engage DG and glial populations

Cell–cell interactions were inferred using CellChat. As sender cells, EAM were predicted to exhibit stronger outgoing interactions than microglia_0 in both control and epilepsy mice (Figure 8A,B). The interaction from EAM to DG is particularly strong (Figure 8A) and involves the Cadm1 pathway and several glutamate-signaling pathways (Figure 8D). The Igf1 signaling pathway was identified as a significant interaction from EAM to several glial and neuronal cell types (Figure 8B,D).

**Figure 8.**
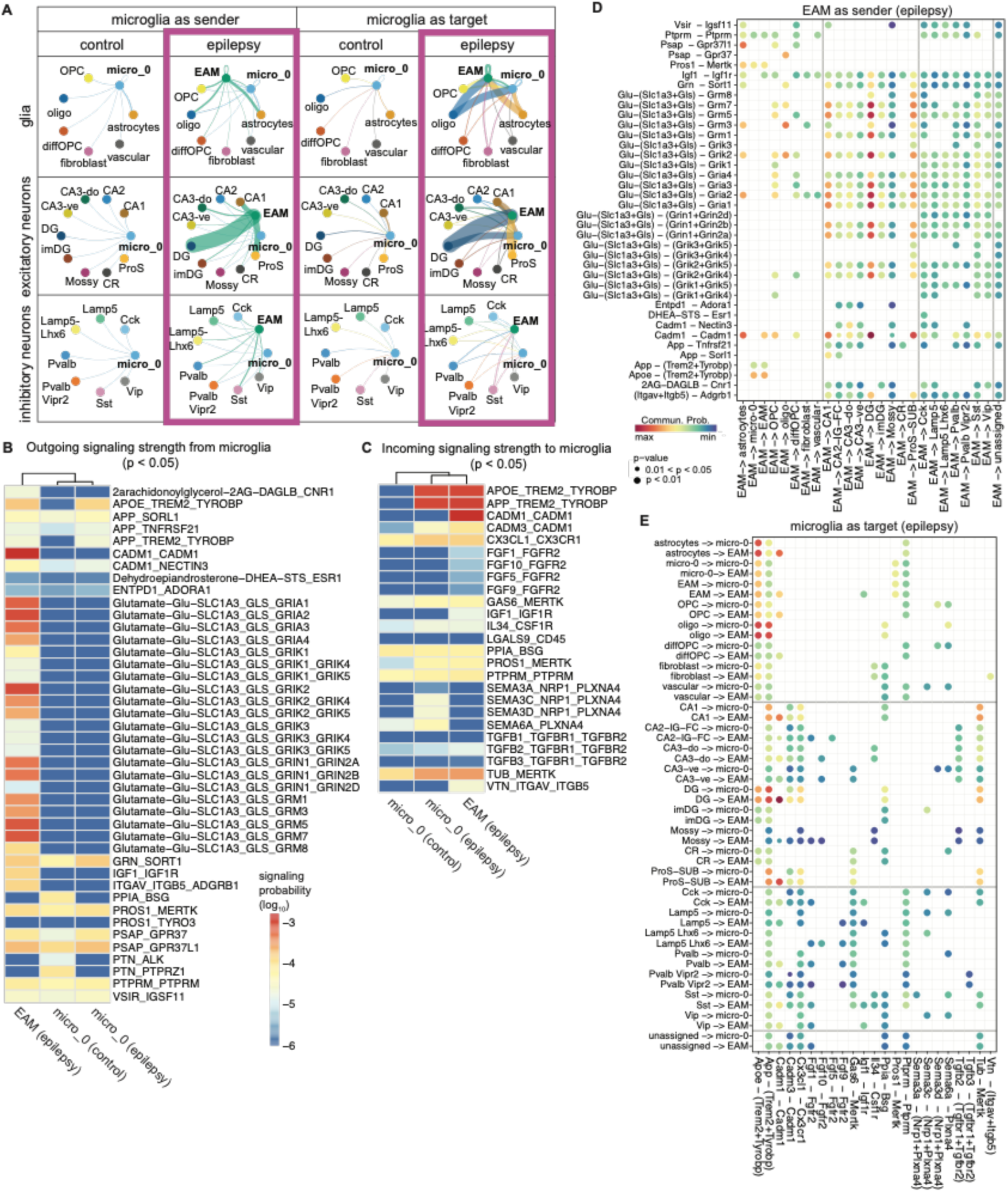
Predicted interactions between microglia and other cell types. (A) Outgoing and incoming microglial interactions in control and epilepsy samples. Edges are colored based on the “sender” cell type and thickness represents interaction weight. The interaction names and probabilities for each microglia cluster are delineated in B and C. The interaction names, partners, and probabilities of the microglia clusters in epilepsy (purple boxes) are delineated in part D and E. (B) Comparison of the outgoing interaction probabilities from the microglia subclusters in epilepsy and control. (C) Comparison of the incoming interaction strengths to the microglia subclusters in epilepsy and control. (D) Interaction names of outgoing EAM interactions in epilepsy and their probabilities. (E) Interaction names of interactions incoming to EAM and microglia_0 in epilepsy and their probabilities. OPC, oligodendrocyte progenitor cell; diffOPC, differentiating oligodendrocyte progenitor cell; DG, dentate granule cells; imDG, immature dentate granule cells; CR, Cajal-Retzius cells; ProS, pro-subiculum - subiculum; EAM, epilepsy-associated microglia. [High-resolution image available as separate file.]

As target cell types, microglia_0 and EAM in epilepsy are both predicted to have stronger interactions compared to microglia_0 in control (Figure 8A,C). The main interaction partner of EAM is again DG cells, and to a lesser extent oligodendrocytes and astrocytes (Figure 8A,E). All three cell types are predicted to have high probabilities of interacting through the Trem2/Tyrobp pathway (Figure 8E), which has previously been found to play a role in the formation of Alzheimer’s disease-associated microglia^23^. The Cadm1 pathway was also predicted to have moderately high interaction probabilities, with astrocytes and several excitatory neurons (CA1, DG, and ProS-SUB) as senders (Figure 8C,E). Other notable predicted sender pathways targeting microglia are Tub-MertK from excitatory neurons and Cx3cl1-Cx3cr1 from both excitatory and inhibitory neurons (Figure 8E). These pathways may contribute to driving and/or maintaining microglial activation in epilepsy. Whether these predicted interactions occur in vivo remains to be validated and will likely depend on the spatial proximity of the interacting cell types.

## Discussion

Our snRNAseq dataset, generated without cell sorting or enrichment, provides a comprehensive overview of cell type-specific perturbations during early stages of epileptogenesis in the mouse hippocampus. The dataset we present here captures the loss of mossy cells and specific subclusters of Cck and Lamp5-Lhx6 interneurons, as well as expansion of a mature DG cell subcluster, neuroblasts, and immature DG cells. We also detected a relative increase in CR cells, consistent with reports of their enhanced persistence in epilepsy^13^. In glial cells, distinct astrocytes and microglia subclusters are increased. Most notably, a microglia subcluster we term EAM is markedly enriched in epilepsy.

Our analyses suggest that microglia play a central role during epileptogenesis. Microglia abundance strongly covaries with shifts in other cell types, and predicted cell-cell interaction patterns are markedly altered in epilepsy. The identification of EAM and marker genes (*Igf1* and *Myo1e*) allows us to examine the role of microglia further. In ISH-FISH studies, EAM exhibited an amoeboid morphology and could be seen in clumps around pyramidal and granule cell body layers of epilepsy samples. The clumps are reminiscent of Alzheimer’s disease-associated microglia clustering around A𝛽 plaques and microglia nodules seen in viral infections and autoimmune diseases^18,24,25^. EAM within clumps exhibit higher levels of *Igf1* and *Myo1e* than isolated EAM, suggesting elevated EAM activity in regions with clumps. Studies on the role of Myo1e in peripheral macrophages suggest that elevated *Myo1e* may reflect enhanced migration or phagocytosis^26,27^. Microglia-derived Igf1 has been shown to regulate neurogenesis during development and may perform similar functions in adulthood^28,29^. In a disease context, activation of the Igf1 pathway in TBI models can be neuroprotective or increase neurogenesis^30,31^.

Both the snRNAseq data and IHC-FISH studies demonstrate inter-sample variability in microgliosis and EAM abundance in epilepsy tissue. Our ANCOVA and correlational studies suggest that the seizure-free duration of animals prior to tissue collection partially accounts for the variability. One interpretation of these findings is that spontaneous seizures acutely trigger these changes in microglia. This explanation would be consistent with prior studies showing that provoked SE leads to microgliosis and activation of microglia^32^. However, in our study, seizure-free duration is confounded by epilepsy severity, which affects seizure frequency. Because each animal contributed only a single terminal measurement, we cannot exclude that animals with more severe epilepsy (higher seizure burden) have chronically elevated microglial density. Hence, further studies are needed to determine whether the microglia response is due to recent spontaneous seizures, epilepsy severity, or both.

In our follow-up studies of EAM, we found that transcriptomic identity does not fully predict functional state. While a subset of microglia in epilepsy samples cluster with “homeostatic” populations in control microglia (i.e. microglia_0), they may be functionally more similar to EAM. Histological analyses indicate that microglia_0 in epilepsy (identified as *Myo1e-; Igf1-*) exhibit morphology that is more similar to EAM (*Myo1e+; Igf1+*) than their counterparts in control samples. Additionally, the incoming cell-cell interaction predictions are more similar between microglia_0 in epilepsy and EAM than with microglia_0 in control. Together, these observations suggest a partial disconnect between transcriptomic clustering and biological function, and underscore the need for caution when inferring microglia roles in disease based solely on transcriptomic profiles.

Although numerous cellular alterations in TLE have been described, teasing out the key drivers of epileptogenesis and their therapeutic relevance has been challenging. Here, we resolve hippocampal cell types at single-nucleus resolution, linking well-established histopathological features of TLE to their underlying transcriptional programs. Together, these findings provide a framework for dissecting the mechanistic contributions of specific cellular populations to epileptogenesis and for evaluating whether targeted modulation of microglial activation can alter disease progression.

## Methods

### Animals

All experiments were conducted in accordance with National Institutes of Health (NIH) guidelines and approved by the Chancellor’s Animal Research Committee at the University of California, Los Angeles. Wild-type C57BL/6J mice were purchased from Charles River Laboratories (strain 027). Single nuclei sequencing studies were performed with male mice only. IHC and FISH studies were performed with male and female mice. The animals were housed on a 12-h/12-h light/dark cycle with access to food and water ad libitum. All experiments were reviewed and overseen by the institutional animal use and care committee at UCLA in accordance with NIH guidelines for the humane treatment of animals.

### Pilocarpine treatment

At 6-weeks old, mice received intraperitoneal (i.p.) injections of scopolamine methyl bromide (1 mg/kg; Sigma-Aldrich, CAS 155-41-9) followed 30 minutes later by a weight-based dose of pilocarpine hydrochloride (weight > 25 g received 250 mg/kg, weight 20-25 g received 275 mg/kg, weight < 20 g received 285 mg/kg; Sigma-Aldrich, CAS 54-71-7).

Littermate control mice received scopolamine only. The mice were subsequently observed visually to identify SE (i.e. continuous, generalized clonus). At 2 hours after the onset of SE, diazepam (10 mg/kg i.p.) was administered to pilocarpine-treated and control mice. The mice were placed on a heat pad, given softened food, and monitored for recovery over 3 days, with 0.5 mL of subcutaneous (s.c.) saline administered daily while appearing lethargic. Pilocarpine-induced SE is expected to produce epilepsy in 100% of mice^33^. All mice that experienced SE had epileptiform discharges on subsequent EEG monitoring (i.e. discharges and/or seizures) and were included in the epilepsy group. Pilocarpine-treated mice that did not enter SE or died during induction were excluded from further study.

### EEG monitoring

Control and pilocarpine-treated mice underwent implantation of stainless-steel screw electrodes 1-week prior to tissue collection. Mice were anesthetized with 1.5–2.0 mL/min of isoflurane in 100% oxygen and fixed in a stereotactic surgery frame. Body temperature was maintained at 37.0-38.0 ^∘^C. Recording electrodes were positioned bilaterally in the parietal skull 1.0 mm posterior and 1.5 mm lateral to bregma. A single ground/reference electrode was positioned in the occipital skull 2.0 mm posterior and 1.0 mm lateral to lambda. The mice received carprofen (5 mg/kg s.c.). Following surgery, the mice were placed on a heat pad and monitored for recovery before being returned to their home cages. EEG data were acquired on an Intan RHD recording system (Intan Technologies, Los Angeles, CA, USA) with wide bandwidth (0.1–7500 Hz) at 1 kHz per channel sampling rate. EEG data were manually reviewed using EDFbrowser to identify seizures and interictal discharges. For the epilepsy group, seizure-free duration was quantified as the time from last seizure to tissue collection or total recording duration if no seizure was present during the recording. For the control group, these values were imputed as zero for ANCOVA.

### Single nucleus isolation and sequencing

At 3- and 6-weeks post-treatment, and following at least 12 hours of seizure-freedom on EEG, pilocarpine-treated and control littermates were processed in batches. Mice were anesthetized with isoflurane and decapitated. Brains were quickly removed and bilateral hippocampi were dissected on ice. Each hippocampus was bisected and the dorsal halves were snap frozen and stored at-80 ^∘^C for nuclei isolation at a later date. Tissue extraction was typically performed at the beginning of the light cycle, although this varied depending on when the last seizure occurred.

Nuclei were extracted as previously described^34^, with modifications. All procedures were conducted on ice or at 4 ^∘^C with precautions to avoid RNAse contamination. Previously frozen pairs of dorsal hippocampal tissue were homogenized using a Dounce homogenizer (Kimble pestle B, DWK Life Sciences 885300-0002) with 1 mL of homogenization buffer (HB): 250 mM sucrose, 25 mM KCl, 5 mM MgCl_#_, 10 mM Tris pH8, 0.1% Nonidet P 40 Substitute (Sigma 74385), 1 μM DL-dithiothreitol (Promega P117B), protease inhibitor (Roche 11697498001), 0.2 U/μL RNase inhibitor (NEB M0314), 1% BSA (VWR EM-2930). The homogenate was kept on ice for 5 min to allow for plasma membrane lysis. The lysate was filtered through a 40 μm Falcon cell strainer (Corning 352340) and centrifuged for 8 min at 600 x g. The pelleted nuclei were resuspended in HB. For purification on a discontinuous iodixanol gradient, the nuclei were combined with an equal volume solution of 50% iodixanol (OptiPrep, Sigma D1556), 250 mM sucrose, 150 mM KCl, 30 mM MgCl_#_, and 60 mM Tris pH8 for final concentration of 25% iodixanol. This mixture was layered over an equal volume of 29% iodixanol and centrifuged at 10,000 x g for 20 minutes. The pelleted nuclei were washed three times in 1 % BSA/PBS solution and passed through a 40 μm filter tip (Sigma BAH136800040). The nuclei were counted and resuspended to 1000 nuclei/μL and submitted to the UCLA Technology Center for Genomics and Bioinformatics for library preparation and sequencing. Single-nuclei libraries were prepared using the Chromium Single Cell 3’ Reagent Kit v3 (10x Genomics) according to the manufacturer’s protocol.

Paired-end sequencing (50 base pair reads) was performed on an Illumina NovaSeq 6000 using the S2 reagent kit. A total of 31 mice were sequenced. One control sample was not sequenced due to too few nuclei extracted.

### Sequencing data processing

Gene-count matrices were generated using the CellRanger v7.1.0 (10x Genomics)’count’ command with the GRCm39 (mm10) mouse transcriptome as reference. Ambient RNA contamination was estimated per sample with SoupX (v1.6.2) autoEstCont and removed with adjustCounts(roundToInt = TRUE) to produce integer, decontaminated UMI counts suitable for downstream methods^35^. Doublet detection was performed with scDblFinder and cells labeled “doublet” were removed^36^. Per-cell QC metrics were determined: total unique molecular identifiers (UMIs) as nCount_RNA, detected genes as nFeature_RNA, and mitochondrial fraction was computed for genes matching “^mt-”. Sample annotations were added to metadata (sample_ID, batch, state, weeks). Additional quality control filtering was applied to exclude nuclei with <500 UMIs, <200 gene counts and>5% mitochondrial gene content. Normalization, scaling, and principal component analysis were performed with Seurat (v5.0.1)^37,38^. Integration was performed using Harmony (v1.2.0) with batch as a covariate^39^. Dimensionality reduction was performed using the’RunUMAP’ command.

### Cell type annotation

Clusters were annotated manually using canonical and more recently identified marker genes^40^. For further classification of excitatory neuron subtypes, the Seurat TransferData function was used with the Allen Institute hippocampal dataset (NCBI GEO GSE185862)^10^. Within the glutamatergic neuron class, only clusters with >50% of cells with prediction scores >0.5 were included for analysis, which led to the exclusion of the “L4/5 IT CTX” subclass. The Hochgerner et al. dentate gyrus dataset was used to further annotate imDG, DG, and astrocyte clusters (NCBI GEO GSE104323)^11^.

To explore whether peripheral immune cells were present in the microglia cluster, the Seurat TransferData function was applied to microglia using several reference datasets downloaded from the Brain Immune Atlas^16,41,42^. SingleR was also used with the “ImmGenData” and “mouse_rnaseq” datasets through celldex^43–45^.

### Cell-type proportion comparisons and network inference

Cell type abundance was studied using the proportion of each cell type within a sample, i.e. cell counts were normalized to the total number of cells per sample such that the proportions within a sample sum to 1. To account for the constraints of compositional data, analyses were performed on zero-imputed, transformed proportions. To assess the effects of treatment and time on cellular composition, we performed a multivariate analysis of variance with treatment (pilocarpine vs. control) and timepoint (3 and 6 weeks post-SE) as fixed factors and included their interaction. The analysis included 22 cell types as dependent variables and was conducted on additive log ratio transformed proportions.

Statistical significance was evaluated using Pillai’s trace. To determine which cell types or subclusters changed in abundance between control and epilepsy, the propeller.ttest method from the speckle R package was applied to arcsin square root transformed cell type proportions^46^. Weeks post-SE was included as a covariate.

Measures of proportionality were determined using the R package propr (v.5.1.8)^47^. Proportionality (𝜌) was computed per condition on centered log-ratio (CLR) transformed proportions using propr (metric = “rho”, ivar = “clr”). FDR was permuted using updateCutoffs (tails = “both”). Differential proportionality (epilepsy − control) was computed using propd. We focused on relationships that shift, i.e. disjointed proportionality (𝜃_*d*), and computed an exact F-test with updateF.

To assess coordinated variation among cell-type proportions, we analyzed CLR–transformed compositional data using a graphical lasso approach. CLR-transformed matrices were normalized and used to estimate sparse inverse covariance matrices with the huge R package (v1.3.5; method = “glasso”)^48^. The regularization parameter was selected using the Stability Approach to Regularization Selection to balance network stability and sparsity, and this parameter was applied uniformly to both control and epilepsy groups to ensure comparable network density and facilitate direct comparison. Nonzero entries in the resulting inverse covariance matrices were interpreted as partial correlations between cell types, representing conditional dependencies after accounting for all other cell types. Differential coupling between conditions was quantified as the difference in partial correlation values between epilepsy and control networks (Δ𝜌).

Adjacency matrices derived from these networks were used to compute graph-theoretical metrics with the igraph package (v2.1.4).

### Differential gene expression analysis

Differential expression testing was performed on pseudobulked data with DESeq2^49^. Raw counts were aggregated at the sample level within each cell type, thereby preserving biological replication and avoiding pseudoreplication. To assess disease progression, testing was performed within epilepsy samples only to compare 3- and 6-week post-SE time points. For epilepsy versus control comparisons, pseudobulk profiles were pooled across both time points. Time-dependent epilepsy effects were evaluated using an interaction model (∼weeks + state + weeks:state) to test whether epilepsy-associated expression changes differed between time points. Genes with <10 counts across samples and clusters with <2 samples per group were excluded. Differential expression was reported as log_#_ fold changes. P-values were adjusted for multiple testing using the Benjamini–Hochberg method.

### Trajectory inference

Cells in the dentate granule cell lineage were merged: RGL cells (i.e. astrocyte subcluster 3) and the DG and imDG clusters. To quantify cluster-cluster connectivity, we used partition-based graph abstraction (PAGA) as implemented in Scanpy (v1.11.5)^50^. Dimensionality reduction was performed using TruncatedSVD. A neighborhood graph was constructed using sc.pp.neighbors (n_neighbors=15). Connectivity scores were computed with sc.tl.paga with the original cluster assignments as the grouping variable.

### Comparison to other disease models

The TBI and AD datasets were downloaded from NCBI GEO (accessions GSE269748 and GSE98969)^17,18^. From the TBI dataset, a Seurat object was created using counts from the 7-day post-CCI peri-contusional tissue and 24-hour naive samples. From the AD dataset, a Seurat object was created using counts from cortex of 6-month 5xFAD and control samples. For each dataset separately, microglia were identified and re-clustered. A distinct microglia subcluster present only in disease samples could be identified in both, in addition to a larger subcluster that was present in control and disease samples. Counts were aggregated for subcluster within each sample, and DESeq2 was used for differential testing. Significant DEGs (adjusted p <0.05) from all three models were compared to identify overlapping DEGs. SuperExactTest (v.1.1.0) was used to calculate significance of the overlap using the union of the DEGs (n = 2865) as background^51^.

### Tissue processing for immunohistochemistry and in situ hybridization

Following at least 12 hours of seizure-freedom on EEG, the mice were anesthetized with isoflurane and trans-cardially perfused with PBS followed by 4% PFA. Brains were quickly removed and fixed in 4% PFA overnight at 4 ^∘^C before paraffin embedding.

For immunohistochemistry, 20 μ m sections were cut on a microtome. Antigen retrieval was performed by boiling slides in sodium citrate (pH 6.0) for 10 minutes. The sections were incubated overnight at 4 ^∘^C in primary antibodies against IBA1 (Thermo Fisher Scientific #PA5-18039, RRID:AB_10982846) and MYO1E (Abcam #ab231789, RRID:AB_3720973). Secondary antibodies (Thermo Fisher Scientific #A32794, RRID:AB_2762834; Thermo Fisher Scientific #A11055, RRID:AB_2534102) were incubated at 1:1000 together with DAPI (Invitrogen #D1306) for 1 hour at room temperature.

Dual IHC-FISH was performed with the RNAscope Multiplex Fluorescent Reagent Kit v2 (ACD #323100) on 5 μ m sections. RNAscope probes against Myo1e (ACD #1238501-C2) and Igf1 (ACD #443901) were applied according to the manufacturer’s protocol with the following modifications: epitope retrieval was performed for 10 minutes and Protease Plus treatment was performed for 5 minutes. TSA Vivid fluorophores 570 and 650(ACD #323272 and #323273) were applied at 1:3000. IHC was performed following completion of the RNAscope signal amplification steps, as previously described^52^. Primary antibody against IBA1 (Abcam #ab178846, RRID:AB_2636859; 1:200) was incubated overnight at 4 ^∘^C, followed by secondary antibody (Thermo Fisher Scientific # A-21206, RRID:AB_2535792) and DAPI for 1 hour at room temperature.

### Image acquisition

Confocal fluorescence imaging was performed on a Zeiss LSM 800 laser-scanning confocal microscope mounted on an Axio Observer.Z1/7 inverted microscope using Plan-Apochromat 20×/0.8 NA air (for IHC only) or 40×/1.4 NA oil-immersion (for dual IHC-FISH) objectives. Fluorophores included DAPI, Alexa Fluor 488/555, Cy3 (TSA 570), and Cy5 (TSA 650), acquired using standard excitation and emission settings. Tile images of the hippocampus were acquired for quantification of microglia density and MYO1E. Z-stacks were acquired with 1.0 µm Z-steps over ∼20 μ m for microglia morphology. Stitching and maximal intensity projections were processed using ZEN 2.6 (blue edition).

### Quantification of microglia density and mRNA puncta

QuPath was used for quantification of microglia and mRNA puncta^53^. Regions of interest (ROIs) were drawn around the hippocampus, CA1, CA2/3, and dentate gyrus. Boundaries of each region were approximated based on the DAPI counterstain, with the CA1c/CA2 boundary determined by where the stratum pyramidale became less compact. Cell segmentation was performed with DAPI, then classified as microglia using a threshold based on the mean IBA1 signal. IBA1^!^ microglia were counted within each ROI and normalized to the ROI area to obtain total microglia density (cells/mm^#^). *Igf1* and *Myo1e* puncta were identified using the “subcellular detection” function, with clusters included for spot count estimation. IBA1^!^ cells that contained ≥ 2 + *Igf1* and ≥ 2 + *Myo1e* puncta were counted as EAM and similarly normalized to ROI area to obtain EAM density (cells/mm^#^).

To assess whether disease state (epilepsy vs. control) was associated with microglial abundance while accounting for potential confounders, we performed an analysis of covariance (ANCOVA) using ordinary least squares regression. Disease state was included as the primary factor, with sex, weeks post–status epilepticus (SE), and seizure-free duration included as covariates. Type II sums of squares were used to obtain 𝐹 statistics and corresponding 𝑝 values for each term.

To examine differences in *Igf1* and *Myo1e* expression levels in greater detail, we calculated histology scores (H-scores) for each ROI^54^. We considered a group of microglia as a clump if there were 4 or more DAPI+ nuclei enclosed by IBA signal with indistinguishable cell boundaries between them. The mRNA puncta content of each cell was binned as follows: 1+ (1 to 3 copies/cell), 2+ (4 to 9 copies/cell), 3+ (10 to 15 copies/cell), and 4+ (>15 copies/cell). The H-score was calculated as (1 × percent of 1+ microglia) + (2 × percent of 2+ microglia) + (3 × percent of 3+ microglia) + (4 × percent of 4+ microglia). Percentages were calculated as a proportion of total microglia. Scores may range between 0-400.

Sex differences in total microglia and EAM density were assessed within the epilepsy group using ANCOVA. Sex (male vs. female) was included as the primary factor, with weeks post-SE and seizure-free duration included as continuous covariates. Statistical significance was defined as 𝑝 < 0.05, and all tests were two-tailed.

### Quantification of microglia morphology

Z-stack images were acquired as described above. For each sample, 2-3 fields of view were taken from each of the CA1, CA2/3, and DG subfields. The fields were selected to contain at least one microglia. Regions containing large numbers of microglia with indistinguishable boundaries (i.e. clumps) were avoided. The acquired images were pre-processed with Fiji (ImageJ, NIH) using the Enhance Contrast and Gaussian Blur functions to isolate the IBA1 signal. Then, 3DMorph was used in interactive mode to skeletonize the signal and calculate the ramification indices (RI; territorial volume divided by the cell volume)^55^. RIs were summarized at the sample level by averaging the RI of all cells within each sample. To determine the *Igf1* and *Myo1e* content of the microglia, maximum intensity projections of the Z-stack images were analyzed with QuPath as described above. Two-sided Mann-Whitney tests were used for pairwise comparisons performed on per-sample mean RI and Holm adjustment was applied.

### Electron microscopy

Brains of animals from control and pilocarpine treated animals (n = 3 per condition) were processed at 6-weeks post-treatment. Following at least 12 hours of seizure-freedom on EEG, mice were deeply anesthetized with isoflurane and were perfused transcardially with a mixture of 4 % paraformaldehyde (PFA) and 0.1 % glutaraldehyde (GA) in 0.1 M phosphate buffer (PB, pH 7.4). Brains were removed and post-fixed in 4% PFA in 0.1 M PB overnight at 4 ^∘^ C. 70 𝜇m thick sections were cut with a Leica VT 1000S vibratome (Leica, Germany). Floating sections were treated for 30 minutes in 1 % NaBH4 in 0.1M PB to quench free aldehyde groups followed by 10 minute incubation in 3 % hydrogen peroxide to eliminate endogenous peroxidase activity. The sections were incubated in 20 % Normal Donkey Serum (NDS) for 30 min to suppress nonspecific binding and then incubated for 12 hr in the primary antibody (Wako #018-28523, RRID:AB_2936184; 1:250), along with 2 % NDS. After rinses in PBS, sections were incubated in biotinylated donkey-anti rabbit IgG (Jackson, Germany) secondary antibody for 2 hrs. After several washes in PB, sections were then incubated in Extravidin peroxidase (1:5000; Sigma, Germany) for 20 min. The immunopositive structures were visualized with 3,3’-diaminobenzidine tetrahydrocholoride (DAB, Sigma, Germany). Sections were processed as described above in control experiments, omitting primary antibody from the incubation solution. In such sections, no immunostaining was observed.

The free-floating sections for electron microscopy were postfixed with 1% OsO4, dehydrated in ascending ethanol series and embedded in epoxy resin (Durcupan; Sigma, Germany) within Aclar sheets (EMS, Hatfield, PA, USA). Uniform rectangular samples were cut from the dorsal hippocampus, at approx.-2.00 to-2.45 mm coronal level from the Bregma under a Leica S6E dissecting microscope and mounted on plastic blocks. 60 nm ultrathin sections were cut on a Reichert ultramicrotome, mounted on 300 mesh copper grids, contrasted with lead citrate (Ultrostain II, Leica, Germany) and examined with a JEM-1011 transmission electron microscope (JEOL, Tokyo, Japan) equipped with a Mega-View-III digital camera and a Soft Imaging System (SIS, Münster, Germany) for the acquisition of the electron micrographs. Five to ten sections were analyzed per block, and two blocks per animal were used to collect micrographs. Sample areas from CA1 strata pyramidale and radiatum of the dorsal hippocampus were chosen in a pseudo-random fashion (at least 50 µm^#^ per animal) on which IBA-1 peroxidase-immunoreactive profiles (cell bodies and glial processes) were identified and photographed. Identified profiles and their intracellular compartment dimensions (vesicles, mitochondria) were measured using the engine provided by NIH ImageJ v1.8.0_345; data were compiled using Excel (Microsoft) software. Statistical significance was determined by Student’s t-test, with a P <0.05 considered statistically significant. Data collection and quantification was performed blindly, to eliminate bias.

### Cell-cell communication inference

Potential cell interactions were predicted from the snRNAseq data using CellChat (v2.2.0) and the built-in mouse interaction database (CellChatDB.mouse) with the standard pipeline: subsetData, identifyOverExpressedGenes, identifyOverExpressedInteractions, computeCommunProb(population.size = TRUE), filterCommunication, and aggregateNet^56^.

For condition-level inference, the dataset was split by disease state (control vs. epilepsy) and CellChat was run separately for each subset. Network centrality was computed and plots were generated with netAnalysis_signalingRole_scatter, netVisual_circle, and netVisual_bubble. Heatmaps were generated using pheatmap (v1.0.13).

For sample-level inference, CellChat was run independently for each sample. For each sample, total outgoing and incoming interaction strength (weight) and interaction counts were summed per cell type. To compare epilepsy vs. control while accounting for batch, analyses were restricted to batches containing at least one epilepsy and one control sample. Within each batch and cell type, epilepsy and control means were computed for each metric and expressed as log_#_(epilepsy/control) ratios; non-finite log_#_ ratios were excluded. Batch-level log_#_ ratios were summarized for each cell type (mean ± SD) and tested against zero using two-tailed one-sample 𝑡 tests, with Benjamini-Hochberg correction across cell types.

## Acknowledgements

We thank Barbara Nagy, Renáta Pop and Eszter Daniló for their excellent technical assistance. We thank Anatol Bragin for help with building our EEG system.

## Bibliography

1. Engel, J. & Pitkänen, A. Biomarkers for epileptogenesis and its treatment. Neuropharmacology 167, 107735 (2020).

2. Löscher, W., Hirsch, L. J. & Schmidt, D. The enigma of the latent period in the development of symptomatic acquired epilepsy — Traditional view versus new concepts. Epilepsy & Behavior 52, 78–92 (2015).

3. Engel, Jr., Jerome. Epileptogenesis. in Seizures and epilepsy 296–319 (Oxford University Press, New York, 2013).

4. Houser, C. R. Neuronal loss and synaptic reorganization in temporal lobe epilepsy: Jasper’s basic mechanisms of the epilepsies. Third edition. Advances in neurology 79, 743–761 (1999).

5. Chauvière, L. et al. Early Deficits in Spatial Memory and Theta Rhythm in Experimental Temporal Lobe Epilepsy. The Journal of Neuroscience 29, 5402–5410 (2009).

6. Lenck-Santini, P.-P. & Holmes, G. L. Altered phase precession and compression of temporal sequences by place cells in epileptic rats. The Journal of neuroscience: the official journal of the Society for Neuroscience 28, 5053–62 (2008).

7. Helmstaedter, C. Effects of chronic epilepsy on declarative memory systems. In Progress in Brain Research vol. 135 439–453 (Elsevier, 2002).

8. Hansen, K. F., Sakamoto, K., Pelz, C., Impey, S. & Obrietan, K. Profiling status epilepticus-induced changes in hippocampal RNA expression using high-throughput RNA sequencing. Scientific Reports 4, 6930 (2014).

9. Shuman, T. et al. Breakdown of spatial coding and interneuron synchronization in epileptic mice. Nature neuroscience 23, 229–238 (2020).

10. Yao, Z. et al. A taxonomy of transcriptomic cell types across the isocortex and hippocampal formation. Cell 184, 3222–3241.e26 (2021).

11. Hochgerner, H., Zeisel, A., Lönnerberg, P. & Linnarsson, S. Conserved properties of dentate gyrus neurogenesis across postnatal development revealed by single-cell RNA sequencing. Nature Neuroscience 21, 290–299 (2018).

12. Feng, Y. et al. Distinct changes to hippocampal and medial entorhinal circuits emerge across the progression of cognitive deficits in epilepsy. Cell Reports 44, 115131 (2025).

13. Blümcke, I. et al. An increase of hippocampal calretinin-immunoreactive neurons correlates with early febrile seizures in temporal lobe epilepsy. Acta Neuropathologica 97, 31–39 (1999).

14. Torre, E. R., Lothman, E. & Steward, O. Glial response to neuronal activity: GFAP-mRNA and protein levels are transiently increased in the hippocampus after seizures. Brain Research 631, 256–264 (1993).

15. Sano, F. et al. Reactive astrocyte-driven epileptogenesis is induced by microglia initially activated following status epilepticus. JCI Insight (2021) doi:10.1172/jci.insight.135391.

16. Van Hove, H. et al. A single-cell atlas of mouse brain macrophages reveals unique transcriptional identities shaped by ontogeny and tissue environment. Nature Neuroscience 22, 1021–1035 (2019).

17. Jha, R. M. et al. A single-cell atlas deconstructs heterogeneity across multiple models in murine traumatic brain injury and identifies novel cell-specific targets. Neuron 112, 3069–3088.e4 (2024).

18. Keren-Shaul, H. et al. A Unique Microglia Type Associated with Restricting Development of Alzheimer’s Disease. Cell 169, 1276–1290.e17 (2017).

19. Ueno, M. et al. Layer V cortical neurons require microglial support for survival during postnatal development. Nature Neuroscience 16, 543–551 (2013).

20. Girón-Pérez, D. A., Piedra-Quintero, Z. L. & Santos-Argumedo, L. Class I myosins: Highly versatile proteins with specific functions in the immune system. Journal of Leukocyte Biology 105, 973–981 (2019).

21. Gao, C., Jiang, J., Tan, Y. & Chen, S. Microglia in neurodegenerative diseases: Mechanism and potential therapeutic targets. Signal Transduction and Targeted Therapy 8, 359 (2023).

22. Espinoza, K. et al. Dynamic changes in mitochondria support phenotypic flexibility of microglia. Nature Communications 16, 11103 (2025).

23. Deczkowska, A. et al. Disease-Associated Microglia: A Universal Immune Sensor of Neurodegeneration. Cell 173, 1073–1081 (2018).

24. Tröscher, A. R. et al. Microglial nodules provide the environment for pathogenic T cells in human encephalitis. Acta Neuropathologica 137, 619–635 (2019).

25. Schwabenland, M. et al. Analyzing microglial phenotypes across neuropathologies: A practical guide. Acta Neuropathologica 142, 923–936 (2021).

26. Wenzel, J., Ouderkirk, J. L., Krendel, M. & Lang, R. Class I myosin *Myo1e* regulates TLR 4-triggered macrophage spreading, chemokine release, and antigen presentation via MHC class II. European Journal of Immunology 45, 225–237 (2015).

27. Barger, S. R. et al. Membrane-cytoskeletal crosstalk mediated by myosin-I regulates adhesion turnover during phagocytosis. Nature Communications 10, 1249 (2019).

28. Yu, D. et al. Microglia regulate GABAergic neurogenesis in prenatal human brain through IGF1. Nature 646, 676–686 (2025).

29. Ziv, Y. et al. Immune cells contribute to the maintenance of neurogenesis and spatial learning abilities in adulthood. Nature Neuroscience 9, 268–275 (2006).

30. Song, Y. et al. Neuroprotective levels of IGF-1 exacerbate epileptogenesis after brain injury. Scientific Reports 6, 32095 (2016).

31. Littlejohn, E. L. et al. IGF1-Stimulated Posttraumatic Hippocampal Remodeling Is Not Dependent on mTOR. Frontiers in Cell and Developmental Biology 9, 663456 (2021).

32. Wyatt-Johnson, S. K., Herr, S. A. & Brewster, A. L. Status Epilepticus Triggers Time-Dependent Alterations in Microglia Abundance and Morphological Phenotypes in the Hippocampus. Frontiers in Neurology 8, 700 (2017).

33. Löscher, W. The holy grail of epilepsy prevention: Preclinical approaches to antiepileptogenic treatments. Neuropharmacology 167, 107605 (2020).

34. Rexach, J. E. et al. Cross-disorder and disease-specific pathways in dementia revealed by single-cell genomics. Cell 187, 5753–5774.e28 (2024).

35. Young, M. D. & Behjati, S. SoupX removes ambient RNA contamination from droplet-based single-cell RNA sequencing data. GigaScience 9, 1–10 (2020).

36. Germain, P.-L., Lun, A., Macnair, W. & Robinson, M. D. Doublet identification in single-cell sequencing data using scDblFinder. F1000Research 10, 979 (2021).

37. Stuart, T. et al. Comprehensive Integration of Single-Cell Data. Cell 177, 1888–1902.e21 (2019).

38. Hao, Y. et al. Dictionary learning for integrative, multimodal and scalable single-cell analysis. Nature Biotechnology 42, 293–304 (2024).

39. Korsunsky, I. et al. Fast, sensitive and accurate integration of single-cell data with Harmony. Nature Methods 16, 1289–1296 (2019).

40. Cembrowski, M. S., Wang, L., Sugino, K., Shields, B. C. & Spruston, N. Hipposeq: A comprehensive RNA-seq database of gene expression in hippocampal principal neurons. eLife 5, 1–22 (2016).

41. Movahedi Lab, Martens, L. & Kancheva, D. Brain immune atlas. (n.d.).

42. De Vlaminck, K. et al. Differential plasticity and fate of brain-resident and recruited macrophages during the onset and resolution of neuroinflammation. Immunity 55, 2085–2102.e9 (2022).

43. Aran, D. et al. Reference-based analysis of lung single-cell sequencing reveals a transitional profibrotic macrophage. Nature Immunology 20, 163–172 (2019).

44. Benayoun, B. A. et al. Remodeling of epigenome and transcriptome landscapes with aging in mice reveals widespread induction of inflammatory responses. Genome Research 29, 697–709 (2019).

45. The Immunological Genome Project Consortium et al. The Immunological Genome Project: Networks of gene expression in immune cells. Nature Immunology 9, 1091–1094 (2008).

46. Phipson, B., et al. *Propeller:* Testing for differences in cell type proportions in single cell data. Bioinformatics 38, 4720–4726 (2022).

47. Quinn, T. P., Richardson, M. F., Lovell, D. & Crowley, T. M. Propr: An R-package for Identifying Proportionally Abundant Features Using Compositional Data Analysis. Scientific Reports 7, 16252 (2017).

48. Zhao, T., Liu, H., Roeder, K., Lafferty, J. & Wasserman, L. The huge Package for High-dimensional Undirected Graph Estimation in R. Journal of machine learning research: JMLR 13, 1059–1062 (2012).

49. Love, M. I., Huber, W. & Anders, S. Moderated estimation of fold change and dispersion for RNA-seq data with DESeq2. Genome Biology 15, 1–21 (2014).

50. Wolf, F. A. et al. PAGA: Graph abstraction reconciles clustering with trajectory inference through a topology preserving map of single cells. Genome Biology 20, 59 (2019).

51. Wang, M., Zhao, Y. & Zhang, B. Efficient Test and Visualization of Multi-Set Intersections. Scientific Reports 5, 16923 (2015).

52. Nilsson, O. R., Kari, L., Rosenke, R. & Steele-Mortimer, O. Protocol for RNA fluorescence in situ hybridization in mouse meningeal whole mounts. STAR Protocols 3, (2022).

53. Bankhead, P., et al. QuPath: Open source software for digital pathology image analysis. Scientific Reports 7, (2017).

54. Advanced Cell Diagnostics. A Guide for RNAscope® Data Analysis. (2017).

55. York, E. M., LeDue, J. M., Bernier, L.-P. & MacVicar, B. A. 3DMorph Automatic Analysis of Microglial Morphology in Three Dimensions from *Ex Vivo* and *In Vivo* Imaging. eneuro 5, ENEURO.0266–18.2018 (2018).

56. Jin, S. et al. Inference and analysis of cell-cell communication using CellChat. Nature Communications 12, 1088 (2021).

