## Extended Data for "Single-Nucleus Transcriptomics Reveals Cell Type-Specific Remodeling and Epilepsy-Associated Microglia"

### Extended Data - Figures

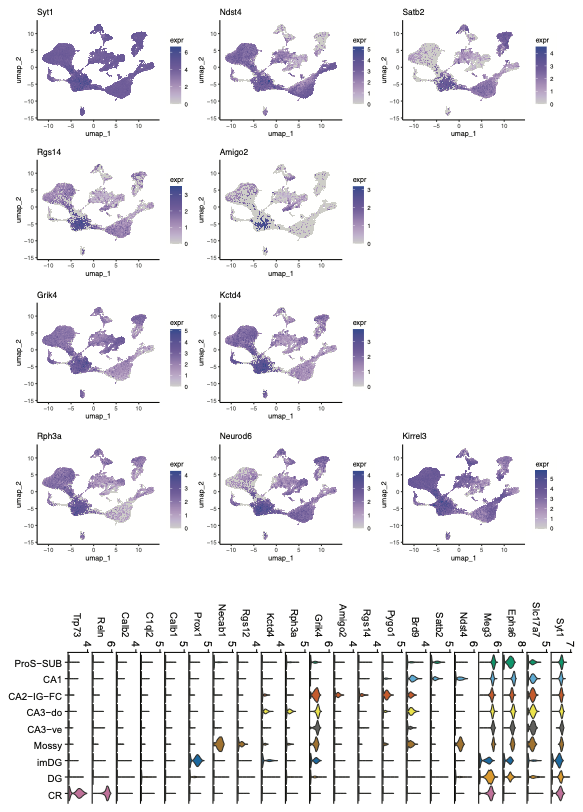

**Figure S1. Expression patterns of marker genes.** Excitatory neuron clusters. Top: Gene expression levels with cells plotted in order of expression. Bottom: Violin plots.

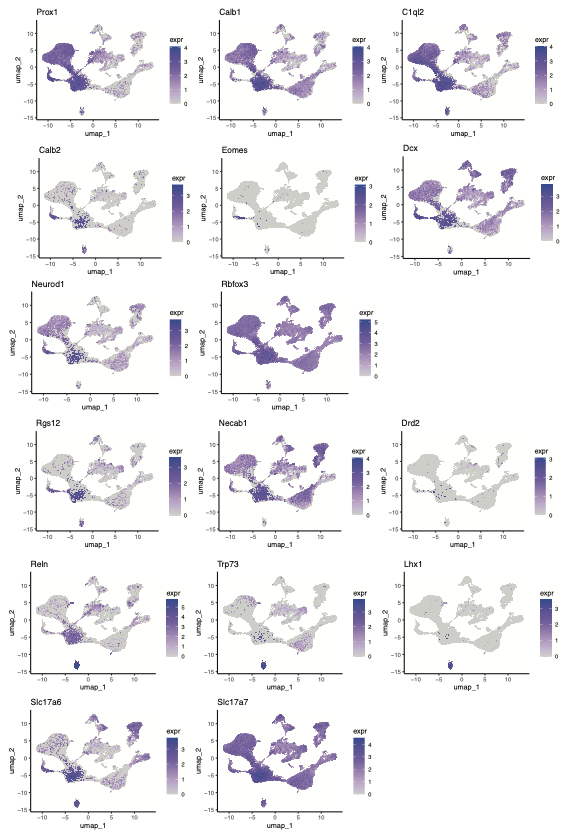

**Figure S1. Expression patterns of marker genes. (continued)** Excitatory neuron clusters. Gene expression levels with cells plotted in order of expression.

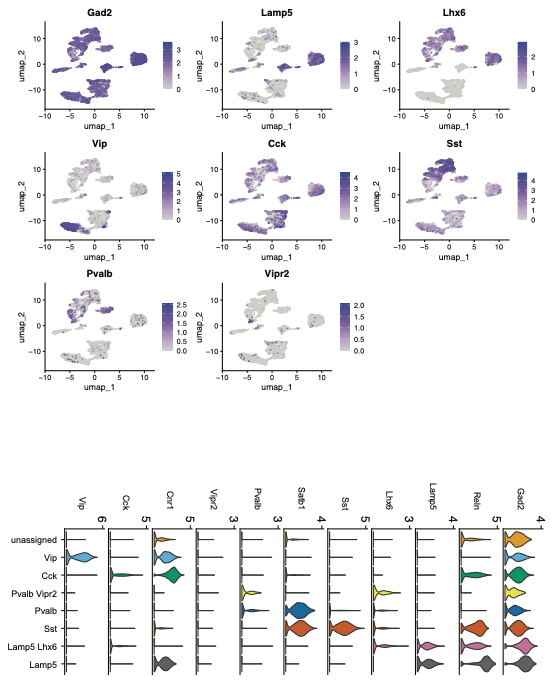

**Figure S1. Expression patterns of marker genes. (continued)** Inhibitory neuron clusters. Top: Gene expression levels with cells plotted in order of expression. Bottom: Violin plots.

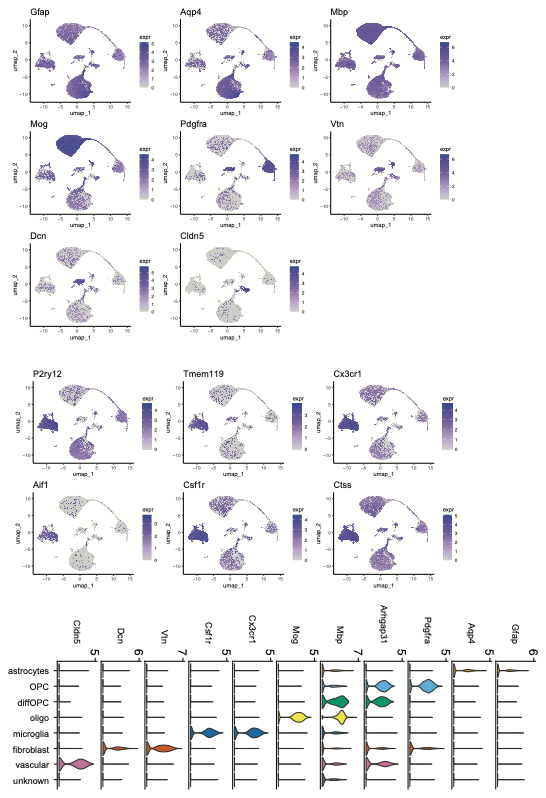

**Figure S1. Expression patterns of marker genes. (continued)** Glial cell clusters. Top: Gene expression levels with cells plotted in order of expression. Bottom: Violin plots.

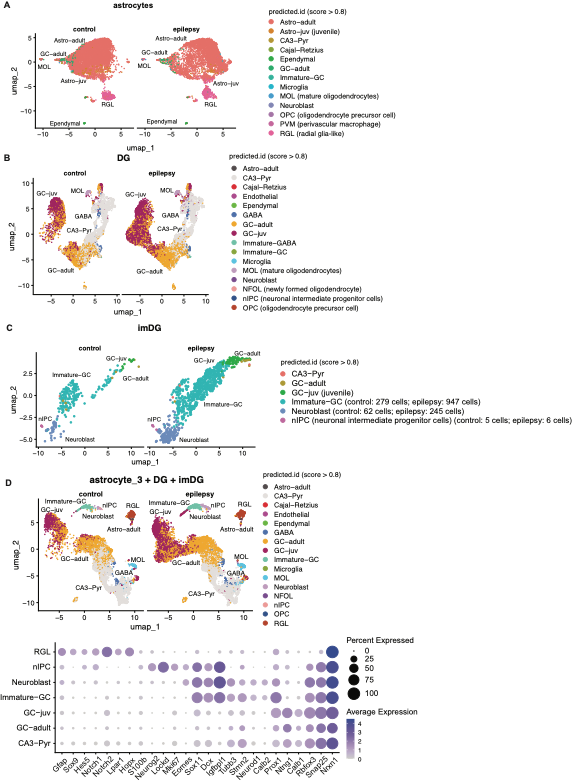

**Figure S2. Dentate granule (DG) cell lineage identified by subclustering and reference-based annotation.** UMAP representations of DG cell lineage-related clusters after subsetting and re-clustering, shown separately for control and epilepsy conditions. Cell identities were assigned using the Hochgerner et al. dentate gyrus dataset as reference^11^. Only annotations with a maximum prediction score >0.8 are displayed. Corresponding subcluster numbers are shown in Figure 4. (A) UMAP of subclusters derived from the astrocyte cluster. Astrocyte subcluster 3 contains cells annotated as radial glia-like (RGL). (B) UMAP of subclusters derived from the DG cluster. DG subclusters 0-2 were annotated as granule cells from juvenile and adult animals. (C) UMAP of subclusters derived from the imDG cluster. Cells in imDG subcluster 0 were annotated as intermediate progenitor cells (nIPCs), neuroblasts, and immature-GC. Cells in imDG subcluster 1 were annotated as immature granule cells (GCs). Cells in imDG subcluster 2 were annotated as a mixture of immature GCs and GCs from juvenile and adult animals. (D) UMAP of merged RGL cells (astrocyte subcluster 3), DG, and imDG clusters after re-clustering. Cells are grouped by their "predicted.id" assigned by TransferData.

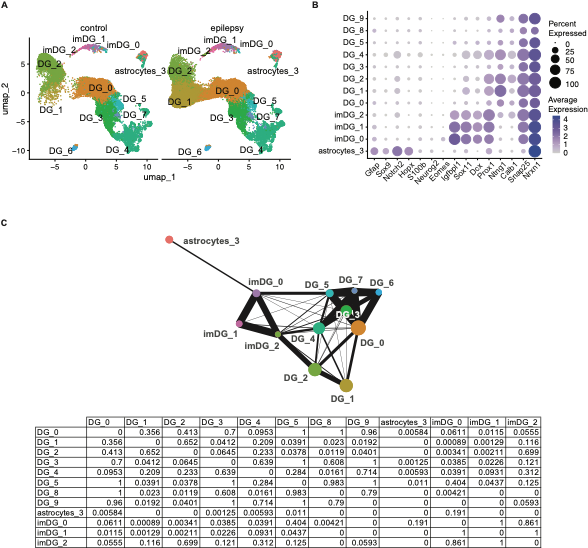

**Figure S3. Trajectory inference of the dentate granule cell lineage.** PAGA analysis identifies DG_1 as a terminal state branching from the DG_2 population. (A) UMAP of merged RGL cells (astrocyte_3), DG, and imDG clusters after re-clustering. Cells are grouped by their original subcluster numbers. (B) Dotplot showing average log-normalized expression of cell type markers. (C) PAGA results. Top: graph where nodes represent clusters and edge weights represent connectivity. Bottom: Matrix showing connectivity scores between clusters. DG_1 has its highest connectivity with DG_2 (0.65).

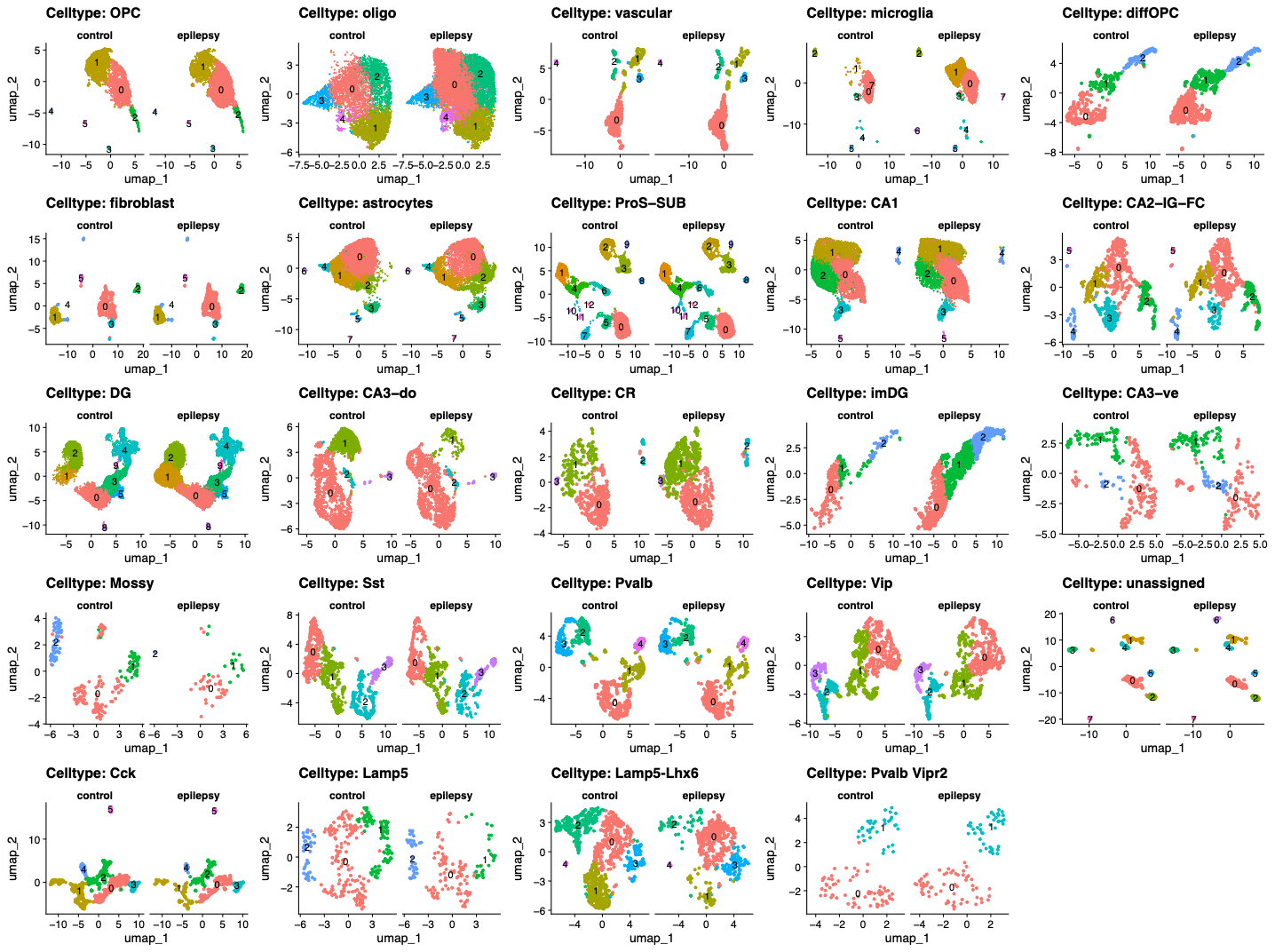

**Figure S4. Re-clustering of cell types identifies subclusters that are gained and lost in epilepsy.** This figure shows all cell types, including those with non-significant differences in proportion between epilepsy and control.

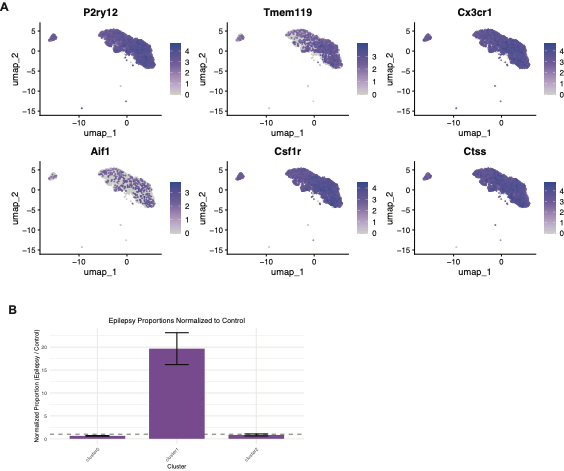

**Figure S5. EAM express canonical microglia markers.** (A) Feature plots showing expression levels of common microglia markers. (B) Proportions of microglia subclusters in epilepsy normalized to control.

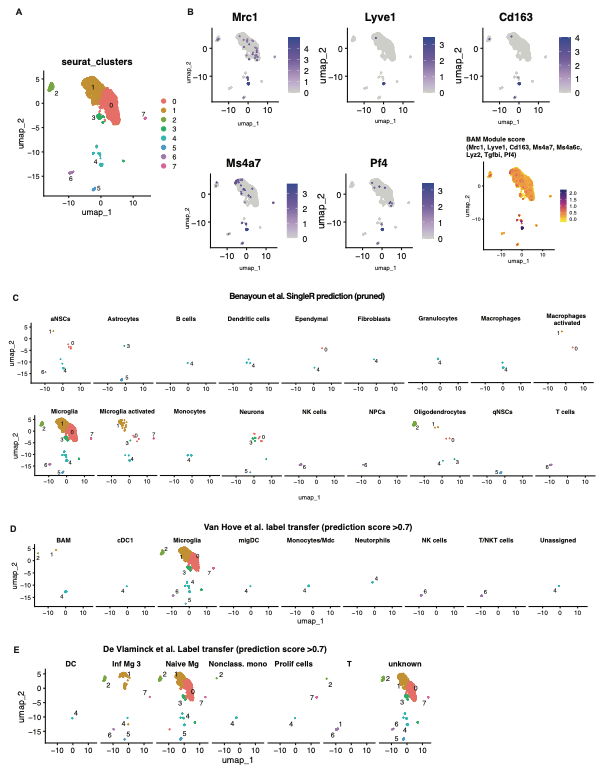

**Figure S6. Immune cell marker expression in the microglia cluster.** Cluster 4 shows high expression of border-associated macrophage (BAM) markers. Automated label transfer using immune cell rich datasets do not identify a robust peripheral immune cell cluster. (A) UMAP of microglia clusters. (B) FeaturePlots showing expression levels of BAM markers in the microglia clusters. The last figure shows the BAM module score, which is derived from the expression levels of the listed genes. (C) The Benayoun et al. dataset was used as reference for automated cell type annotation with SingleR. The resulting classifications are shown, with original cluster numbers preserved. (D) The Van Hove e al. data set was used as reference for automated cell type annotation using the TransferData function in Seurat. Predicted ID scores >0.7 were applied. (E) The De Vlaminck et al. dataset was used as in (D).

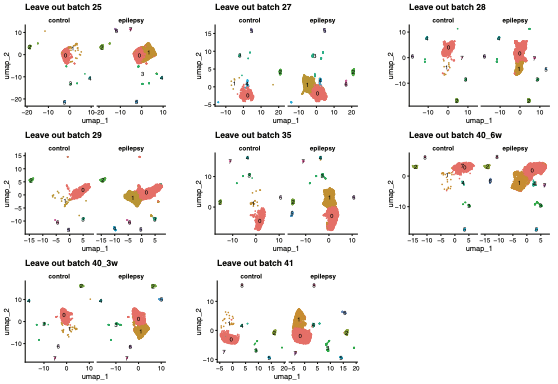

**Figure S7. Leave one out robustness analysis.** UMAP plots from leave-one-out analysis, in which each batch was excluded and the data re-normalized, integrated and recluatered. Across each iteration, an epilepsy-enriched microglia cluster is observed, indicating that the cluster is not driven by a single batch.

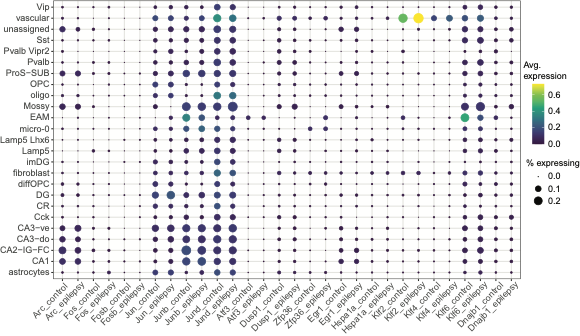

**Figure S8. Immediate early gene expression is low across cell types.** Dot size reflects percentage of cells with detectable expression (>0 raw count), while dot color indicates avearge log-normalized expression from the Seurat "data" slot.

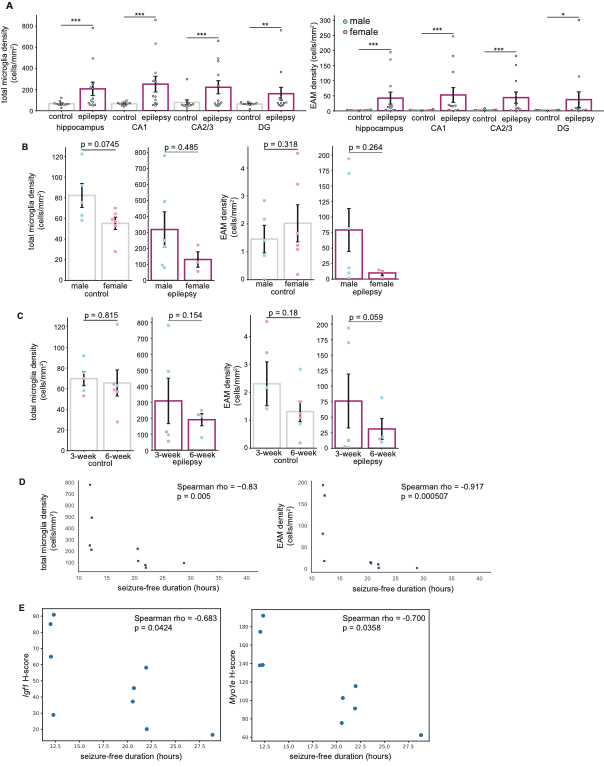

**Figure S9. Using dual IHC-FISH to evaluate covariates that may influence microglia and EAM quantity**. (A) Total microglia and EAM density across hippocampal subregions in control and epilepsy animals. (B) Total microglia and EAM density in male vs. female animals. Control and epilepsy samples shown separately. Each dot represents one mouse. (C) Total microglia and EAM density in animals at the 3- vs. 6-week post-SE timepoints. Control and epilepsy samples shown separately. Each dot represents one mouse. (D) Total microglia and EAM density (quantified by IHC-FISH) plotted against seizure-free duration. Each dot represents one mouse. (E) Igf1 (left) and Myo1e (right) H-scores from hippocampal sections plot against seizure-free duration. Each dot represents one mouse. All error bars show S.E.M.

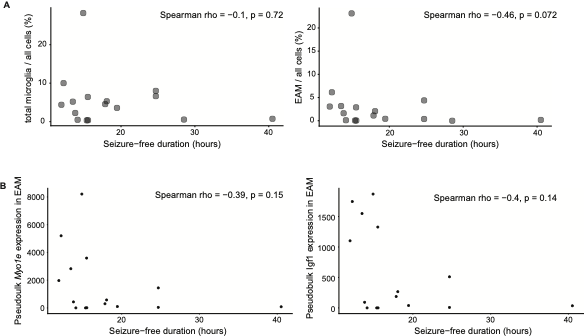

**Figure S10. Using snRNAseq data to evaluate the effects of seizure-free duration on EAM.** (A) Proportion of total microglia (left) and EAM (right) expressed as a percentage of all cells plotted against seizure-free duration. Each dot represents one mouse. (C) Pseudobulked Myo1e (left) and Igf1 (right) expression levels plotted against seizure-free duration. Each dot represents one mouse.

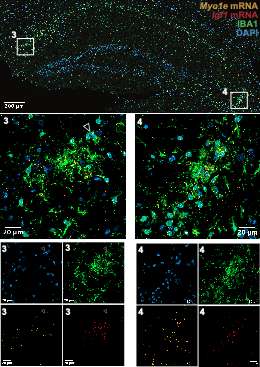

**Figure S11. EAM may be found in clumps around pyramidal cell body layers.** Insets enlarged in panels below and numbered accordingly.

### Extended Data - Tables

**Extended Data Table 1. Differentially expressed genes (DEGs) identified across cell types using DESeq2 on pseudobulked data.** Lists of differentially expressed genes between epilepsy and control (both time points combined) or between 6- and 3-week epilepsy samples was assessed using a pseudobulk DESeq2 framework within each cell type. The last sheet is a list of two significant genes identified by interaction analysis using a DESeq2 design of ~weeks + state + weeks:state

See supplementary file pseudobulk-DEGs.xlsx

**Extended Data Table 2. Cell type proportions.** Cell counts for each cell type normalized to the total number of nuclei sequenced per sample.

See supplementary file celltype-proportions.csv

**Extended Data Table 3. Multivariate Analysis of Variance (MANOVA).** Effects of treatment and timepoint on cell type composition in control vs epilepsy. Multivariate analysis of variance performed on additive log-ratio transformed proportions, with vascular cells as the reference cell type.

| **Effect** | **Df** | **Pillai** | **Approx F** | **Num Df** | **Den Df** | **Pr(**$>$**F)** |
| --- | --- | --- | --- | --- | --- | --- |
| timepoint | 1 | 0.814 | 1.1934 | 22 | 6 | 0.4460 |
| treatment | 1 | 0.989 | 24.3483 | 22 | 6 | **0.0004** |
| timepoint:treatment | 1 | 0.645 | 0.4952 | 22 | 6 | 0.8939 |
| Residuals | 27 |  |  |  |  |  |

**Extended Data Table 4. Comparison of cell type proportions in control vs. epilepsy.** Output from the propeller function applied using arcsin square root transformation and taking timepoint as a covariate. P-values were adjusted for multiple testing using the Benjamini-Hochberg false discovery rate (FDR).

| **Cell Type** | **Control** | **Epilepsy** | **Ratio** | **T-stat** | **P-value** | **FDR** |
| --- | --- | --- | --- | --- | --- | --- |
| Oligo | 0.0745 | 0.1384 | 1.86 | 3.62 | 0.0011 | **0.0174** |
| imDG | 0.0053 | 0.0150 | 2.83 | 3.48 | 0.0016 | **0.0174** |
| Mossy cells | 0.0029 | 0.0010 | 0.35 | -3.35 | 0.0022 | **0.0174** |
| Cck | 0.0162 | 0.0075 | 0.46 | -3.20 | 0.0032 | **0.0193** |
| Microglia | 0.0168 | 0.0621 | 3.70 | 2.85 | 0.0078 | **0.0375** |
| CR | 0.0073 | 0.0111 | 1.51 | 2.76 | 0.0098 | **0.0392** |
| Astrocytes | 0.1402 | 0.0780 | 0.56 | -2.40 | 0.0230 | 0.0700 |
| Lamp5 Lhx6 | 0.0134 | 0.0077 | 0.58 | -2.39 | 0.0233 | 0.0700 |
| diffOPC | 0.0049 | 0.0082 | 1.70 | 1.97 | 0.0587 | 0.1433 |
| Lamp5 | 0.0037 | 0.0020 | 0.55 | -1.96 | 0.0597 | 0.1433 |
| Fibroblasts | 0.0143 | 0.0175 | 1.23 | 1.32 | 0.1977 | 0.4314 |
| CA1 | 0.1367 | 0.1095 | 0.80 | -1.20 | 0.2405 | 0.4632 |
| Pvalb | 0.0104 | 0.0079 | 0.76 | -1.16 | 0.2533 | 0.4632 |
| CA3-do | 0.0256 | 0.0214 | 0.84 | -1.12 | 0.2702 | 0.4632 |
| Sst | 0.0178 | 0.0145 | 0.82 | -0.99 | 0.3324 | 0.5319 |
| Unassigned | 0.0045 | 0.0050 | 1.10 | 0.80 | 0.4290 | 0.6435 |
| Pvalb Vipr2 | 0.0014 | 0.0014 | 1.06 | 0.60 | 0.5502 | 0.7744 |
| Vip | 0.0103 | 0.0086 | 0.83 | -0.56 | 0.5808 | 0.7744 |
| OPCs | 0.0374 | 0.0319 | 0.85 | -0.35 | 0.7264 | 0.9176 |
| CA2-IG-FC | 0.0129 | 0.0139 | 1.08 | 0.18 | 0.8604 | 0.9390 |
| ProS-SUB | 0.1011 | 0.0996 | 0.98 | 0.15 | 0.8799 | 0.9390 |
| Vascular | 0.0075 | 0.0069 | 0.92 | 0.14 | 0.8920 | 0.9390 |
| DG | 0.3311 | 0.3265 | 0.99 | -0.13 | 0.8999 | 0.9390 |
| CA3-ve | 0.0038 | 0.0043 | 1.14 | 0.00 | 0.9994 | 0.9994 |

**Extended Data Table 5. Summary of differential proportionality (**$\Delta\rho=\rho_{epilepsy}-\rho_{control}$**) across cell-class pairs in epilepsy versus control.** For pairs significant by the F-test (FDR $\leq0.05$), the table reports the number of significant pairs ($n_{pairs}$) and aggregate statistics: sum_rho_diff (sum of $\Delta\rho$), sum_pos (sum of positive $\Delta\rho$), sum_neg (sum of negative $\Delta\rho$), and sum_abs (the total magnitude of change), where sum_abs $=\sum_{i} |\Delta\rho_{i}|$ within each class. Positive values indicate stronger proportional co-variation in epilepsy.

| pair_class | n_pairs | sum_rho_diff | sum_pos | sum_neg | sum_abs |
| --- | --- | --- | --- | --- | --- |
| glia-interneuron | 28 | 5.835 | 7.026 | -1.191 | 8.216 |
| excitatory-interneuron | 22 | 3.343 | 5.482 | -2.139 | 7.621 |
| excitatory-glia | 19 | -1.311 | 1.680 | -2.991 | 4.671 |
| glia-glia | 13 | -1.383 | 0.626 | -2.010 | 2.636 |
| excitatory-excitatory | 11 | 0.142 | 0.927 | -0.785 | 1.712 |
| interneuron-interneuron | 9 | -1.203 | 0.139 | -1.342 | 1.481 |
| interneuron-other | 2 | -0.664 | 0.000 | -0.664 | 0.664 |
| excitatory-other | 2 | 0.116 | 0.116 | 0.000 | 0.116 |
| glia-other | 1 | 0.071 | 0.071 | 0.000 | 0.071 |

**Extended Data Table 6. Cell type subcluster comparisons.** See accompanying Excel file. First sheet contains comparisons of subcluster proportions within each cluster. Results are the output from the propeller function applied using arcsin square root transformation and taking timepoint as a covariate. P-values were adjusted for multiple testing across clusters using the Benjamini-Hochberg false discovery rate (FDR). Subsequent sheets show gene enrichment for the indicated subclusters, calculated using DESeq2 on pseudobulked data.

See supplementary file subclusters.xlsx

**Extended Data Table 7. Per-sample cell-cell interactions.** Full per-sample incoming and outgoing interaction weights and counts predicted by CellChat, along with summary statistics (mean, standard deviation, unpaired t-test p-values, and Benjamini–Hochberg adjusted p-values) across batches. See accompanying supplementary Excel file for complete data (first sheet: raw values per sample; subsequent sheets: summaries and statistics). Sample IDs include batch numbers.

See supplementary file cellchat_persample.xlsx

**Extended Data Table 8. DEGs in disease-associated microglia found in EAM, DAM, and TBI.** Full lists of genes differentially expressed in the disease-associated microglial subcluster relative to the homeostatic subcluster shared between control and epilepsy conditions. Differential expression was performed on pseudobulk data using DESeq2 and filtered for adjusted $p<0.05$.

See supplementary file EAM-DAM-TBI-microglia-overlap_genes.xlsx

**Extended Data Table 9. Quantification of total microglia and EAM density by dual IHC-FISH.** Density measurements (cells per mm${}^{2}$) of total microglia and EAM across hippocampal subregions in control and epilepsy animals.

See supplementary file IHC-quantification.csv

**Extended Data Table 10. Analysis of covariance (ANCOVA) for total microglia density.** Statistical testing performed on total microglia density quantified from dual IHC-FISH hippocampal sections. For each region separately, an ANCOVA was fit using: $Y\sim\text{C(state)}+\text{C(sex)}+\text{weeks post-SE}+\text{seizure-free duration}$, where $Y$ is total microglia (cells/mm${}^{2}$), state is a 2-level factor (control, epilepsy), sex is a 2-level factor (female, male), weeks is a continuous predictor, and seizure-free duration is a continuous covariate in days (e.g. 0.5 = 12 h; controls imputed as 0).

| Region | $F_{\text{state}}$ | $p_{\text{state}}$ | $F_{\text{sz-free}}$ | $p_{\text{sz-free}}$ | $p_{\text{weeks}}$ | $p_{\text{sex}}$ | $R^{2}$ |
| --- | --- | --- | --- | --- | --- | --- | --- |
| Hippocampus | 21.59 | $3.16\times{10}^{-4}$ | 14.21 | $1.86\times{10}^{-3}$ | 0.148 | 0.517 | 0.673 |
| CA1 | 29.98 | $6.40\times{10}^{-5}$ | 18.56 | $6.22\times{10}^{-4}$ | 0.122 | 0.496 | 0.743 |
| CA2/3 | 41.37 | $1.13\times{10}^{-5}$ | 28.53 | $8.20\times{10}^{-5}$ | 0.764 | 0.122 | 0.799 |
| DG | 9.84 | $6.79\times{10}^{-3}$ | 6.90 | $1.91\times{10}^{-2}$ | 0.086 | 0.822 | 0.501 |

**Extended Data Table 11. ANCOVA for EAM density.** Statistical testing performed on EAM density quantified from dual IHC-FISH hippocampal sections. For each region separately, an ANCOVA was fit using: $Y\sim\text{C(state)}+\text{C(sex)}+\text{weeks post-SE}+\text{seizure-free duration}$, where $Y$ is EAM density (cells/mm${}^{2}$), state is a 2-level factor (control, epilepsy), sex is a 2-level factor (female, male), weeks is a continuous predictor, and seizure-free duration is a continuous covariate in days (e.g. 0.5 = 12 h; controls imputed as 0).

| Region | $F_{\text{state}}$ | $p_{\text{state}}$ | $F_{\text{sz-free}}$ | $p_{\text{sz-free}}$ | $p_{\text{weeks}}$ | $p_{\text{sex}}$ | $R^{2}$ |
| --- | --- | --- | --- | --- | --- | --- | --- |
| Hippocampus | 26.25 | $1.25\times{10}^{-4}$ | 18.73 | $5.98\times{10}^{-4}$ | 0.076 | 0.577 | 0.705 |
| CA1 | 26.01 | $1.31\times{10}^{-4}$ | 18.12 | $6.89\times{10}^{-4}$ | 0.117 | 0.556 | 0.702 |
| CA2/3 | 36.72 | $2.20\times{10}^{-5}$ | 25.26 | $1.51\times{10}^{-4}$ | 0.800 | 0.336 | 0.766 |
| DG | 8.16 | $1.20\times{10}^{-2}$ | 6.16 | $2.54\times{10}^{-2}$ | 0.109 | 0.809 | 0.450 |
